# Cell type composition drives patient stratification in single-cell RNA-seq cohorts

**DOI:** 10.64898/2026.03.27.714811

**Authors:** Christian Halter, Massimo Andreatta, Santiago J. Carmona

## Abstract

Early transcriptomic studies demonstrated that unsupervised analysis of bulk gene expression can reveal clinically meaningful patient subgroups. Single-cell RNA sequencing (scRNA-seq) provides high-resolution characterization of cellular heterogeneity and therefore enables more refined patient stratification. Several computational approaches have been proposed to summarize single-cell data into sample-level representations for cohort-level exploratory analyses. However, these methods generally do not explicitly account for the compositional nature of cell-type proportions. Based on eleven scRNA-seq cohorts across different biological conditions, we evaluated several state-of-the-art sample representation methods for their ability to recover known biological groupings in an unsupervised setting. Surprisingly, we found that baseline approaches based on cell-type composition and pseudobulk gene expression consistently matched or outperformed more complex methods while requiring orders of magnitude fewer computational resources. In particular, centered log-ratio-transformed cell-type proportions achieved the highest stratification performance and demonstrated robustness to batch effects. The stratification signal was frequently concentrated in a small subset of highly variable cell types, and performance was robust across diverse cell type annotation strategies. Altogether, these results suggest that clinically relevant inter-sample variation in scRNA-seq cohorts is largely driven by differences in cell-type composition. Importantly, compositional representations directly link cohort-level structure to specific cell populations, enabling mechanistic interpretation and facilitating clinical translation. We provide scECODA, an open-source R package for scalable and interpretable cohort-level Exploratory COmpositional Data Analysis of scRNA-seq data, and establish cell-type compositional representations as a powerful and interpretable baseline for unsupervised patient stratification.

## Introduction

For over twenty years, the study of gene expression profiles from bulk transcriptomics has revealed clinically meaningful patient subgroups (Golub et al. 1999, Perou et al. 2000, Sørlie et al. 2001), a paradigm that was subsequently generalized by large-scale efforts such as The Cancer Genome Atlas (Hoadley et al. 2014, 2018). Despite their important insights, these studies relied on bulk RNA profiling approaches, which provide only average molecular profiles of whole-tissue samples. As a consequence, they obscure underlying cellular heterogeneity, including variation in cell-type composition and cell-type-specific gene expression programs, potentially limiting the resolution and interpretability of omics-based patient stratification. The advent of single-cell transcriptomics, which enables molecular profiling at cellular resolution and is increasingly applied to large patient cohorts, provides an opportunity to overcome these limitations. By explicitly capturing fine-grained cell-type composition and cell type-specific gene expression patterns, single-cell RNA sequencing (scRNA-seq) has the potential to enable more refined patient stratification.

From a computational perspective, unsupervised patient stratification involves mapping high-dimensional molecular profiles into a lower-dimensional patient-level representation space, in which biologically meaningful cohort-level structure can be identified, for example through clustering. Compared to bulk profiling, single-cell omics-based patient stratification introduces additional computational complexity, as it requires summarizing information from distributions of single-cell transcriptomes, into patient-level (or more generally, sample-level) representations. Several approaches have been proposed for learning such representations from scRNA-seq data, including matrix- and tensor-factorization methods (MOFA+ (Argelaguet et al. 2020), scITD (Mitchel et al. 2022)), optimal transport and density-based approaches (PILOT (JoodakiMehdi et al. 2023), GloScope (Wang et al. 2024)), and deep generative models (MrVI (Boyeau et al. 2024), scPoli (De Donno et al. 2023)).

Many pathological processes involve the proliferation, depletion, or redistribution of specific cell populations, leading to systematic shifts in cell-type abundances across patients. Cell counts determined by tissue profiling techniques such as scRNA-seq carry only relative information: when the abundance of one cell type changes, the relative abundances of all other cell types necessarily shift. Consequently, cell-type proportions constitute compositional data – they are constrained to sum to one and reside in a subspace known as “simplex” rather than in Euclidean space. Applying standard distance metrics or clustering algorithms to compositional data leads to distortions in inter-sample relationships (Quinn et al. 2018).

Several methods have been developed for comparative analysis of cell-type compositions in scRNA-seq data, commonly referred to as cell type differential abundance analysis (e.g. sccomp (Mangiola et al. 2023), scCODA (Büttner et al. 2021), propeller (Phipson et al. 2022), voomCLR (Assefa, Verbist, and Berge 2024), crumblr (Hoffman and Roussos 2025)). These approaches are designed to test predefined case–control contrasts and assess statistically significant shifts in cell-type proportions. In contrast, compositional analysis in settings where patient groupings are unknown – such as exploratory cohort-level analysis and unsupervised patient stratification – has not been systematically investigated. Existing sample representation methods that rely on cell-type annotations do not explicitly model cell-type proportions as compositional data. As a result, the geometry induced by these representations may fail to capture inter-patient variation when abundance shifts are a dominant driver of biological differences.

Here, we benchmarked state-of-the-art scRNA-seq sample representation methods alongside compositional-aware cell-type proportions embeddings and pseudobulk gene expression representations across 11 patient cohorts comprising 697 samples. We evaluated each method’s ability to recover known patient subgroups in an unsupervised setting. Based on these results, we provide practical recommendations for cohort-level exploratory analysis of scRNA-seq data, considering biological signal recovery, computational efficiency, and interpretability, and establish principled cell type compositional baselines for future sample representation methods.

## Results

### Cell type compositional representation of scRNA-seq data enables highly interpretable cohort-level exploratory analysis

Cohort-level analyses of scRNA-seq data require transforming high-dimensional cell-level molecular profiles into sample-level representations suitable for modeling inter-sample variation. In this study, we propose a simple and interpretable baseline approach for scRNA-seq sample-level representation based on centered log-ratio (CLR)-transformed cell type composition (Figure 1A, see Methods for details). To illustrate how this representation can help reveal the underlying structure of patient cohorts and its relationship to biological variables, we applied it to a large cohort (*n* = 868) of scRNA-seq samples from blood of healthy individuals of known age and cytomegalovirus (CMV) serological status (Gong et al. 2024) (Supplementary table 1).

**Figure 1:**
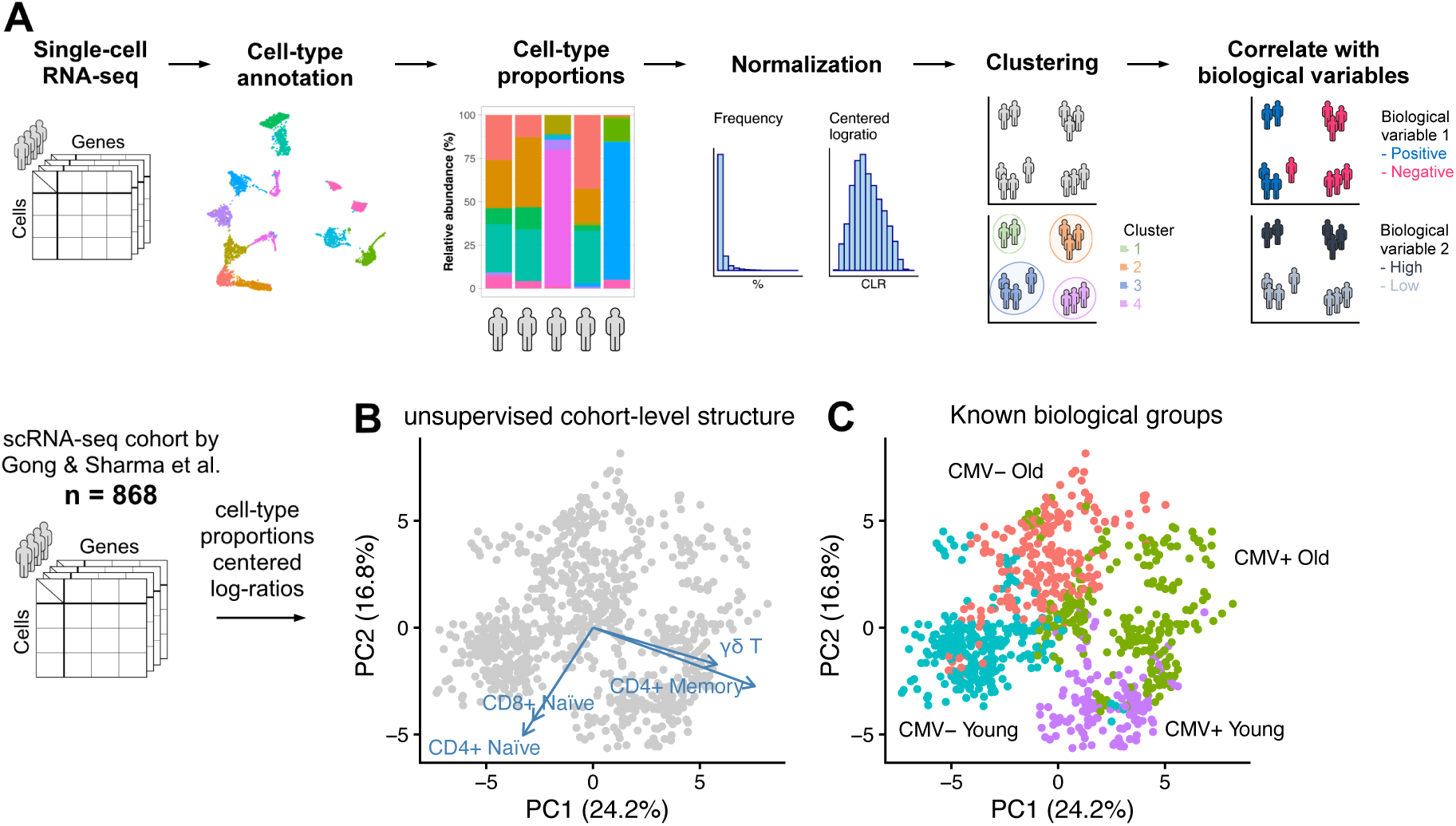
Exploratory COmpositional Data Analysis (ECODA) enables highly interpretable cohort-level exploratory analysis of single-cell transcriptomics data. **A)** Following single-cell RNA-seq data acquisition and cell-type annotation, cell-type compositions are calculated for each sample. Cell-type counts are normalized using the centered log-ratio (CLR) transformation. The resulting sample-level representations are visualized by principal component analysis (PCA), and distinct groups of samples can be identified via unsupervised clustering. **B-C)** Illustrative example of cell-type compositional analysis of a scRNA-seq dataset of blood samples from a cohort of healthy donors of known age group and cytomegalovirus (CMV) serological status. Each point represents an individual donor sample projected onto the first two principal components (PC1 and PC2), which capture the largest axes of variance in cell-type composition across the cohort. **B)** PCA biplot where the arrows represent the top 4 variables (cell types) ranked by their contribution to the two first principal components (see Methods). The arrows point in the direction of increasing centered log-ratio (CLR)-transformed abundance of each cell type, where the CLR-transformation expresses each cell type’s proportion relative to the geometric mean of all cell types within each donor sample. **C)** PCA scatterplot where donor samples are colored by known biological groups.

Principal components analysis (PCA) of CLR-transformed cell-type compositions revealed that the first and second principal components largely captured differences in age and CMV status across individuals (Figure 1B, C). As shown in the PCA biplot (Figure 1B), the features most strongly contributing to these components were specific subtypes of T cells: CMV+ individuals were associated with higher relative abundance of specific subsets of γδ T cells and CD4+ effector memory T cells compared to CMV negative individuals, whereas younger individuals showed higher relative abundance of naïve T cells, compared to older individuals (Figure 1C), in agreement with previous reports (Gong et al. 2024, Pitard et al. 2008, Sylwester et al. 2005, Jia et al. 2023).

This example illustrates that CLR-transformed cell-type proportions can provide informative and highly interpretable sample-level representation of scRNA-seq data, with potential utility for unsupervised patient stratification in cohort-level analyses. We refer to this approach as Exploratory COmpositional Data Analysis (ECODA).

### ECODA outperforms state-of-the-art sample representation methods for unsupervised patient stratification

To evaluate the ability of different scRNA-seq sample representations to recover biologically relevant patient stratifications, we collected scRNA-seq datasets for 11 patient cohorts comprising 697 individuals. Each cohort includes one or more cohort-specific clinical or biological annotations, such as disease labels – spanning malignant, autoimmune, and infectious conditions (e.g. CMV serostatus in Figure 1) – and treatment-related variables, which we used as reference variables for evaluation (Supplementary table 2). These datasets were selected based on two inclusion criteria: i) the presence of at least one well-defined biological condition serving as a ground-truth stratification variable; ii) a cohort size of more than 25 samples with at least 500 cells, to ensure stable estimations.

We conducted a literature survey of scRNA-seq sample-level representation methods and identified approaches based on variational autoencoders (MrVI (Boyeau et al. 2024), scPoli (De Donno et al. 2023)), distribution distances (PILOT (JoodakiMehdi et al. 2023), GloScope, and its variant focused on cell-type proportions GloProp (Wang et al. 2024)) and factor decomposition (MOFA+ (Argelaguet et al. 2020), scITD (Mitchel et al. 2022)) (Supplementary table 3). All methods output sample representations either as sample embeddings or inter-sample dissimilarity matrices. These state-of-the-art methods, together with two baseline sample representation approaches, i) ECODA and ii) whole-sample average gene expression (hereafter, “pseudobulk”), were systematically evaluated for their ability to separate sample groups according to the known biological conditions (see Methods). Separation performance was quantified using three complementary metrics: Adjusted Rand Index (ARI), graph Modularity, and Analysis of Similarities (ANOSIM) (see Methods). The three metrics exhibited strong concordance across all datasets and representation methods (Supplementary figure 1). All tested methods were evaluated across a range of key parameters to ensure that their recommended configurations exhibited stable performance (Supplementary figure 2).

Notably, the patient stratification benchmark revealed that ECODA achieved the highest performance across all three metrics and for the majority of biological conditions (Figure 2A). GloProp and the pseudobulk baseline ranked second and third, respectively, and were followed by GloScope, MOFA, MrVI and scPoli. PILOT and scITD showed the lowest performance. As expected, a negative control compositional baseline with shuffled sample labels (Shuffled_baseline) showed near-zero stratification performance. Two-dimensional sample projections using multi-dimensional scaling (MDS) qualitatively supported the quantitative benchmark results (Figure 2B, Supplementary figure 3 to Supplementary figure 13). For instance, ECODA and GloProp most clearly separated breast cancer patients who responded to cancer immunotherapy from those who did not (Figure 2B).

**Figure 2:**
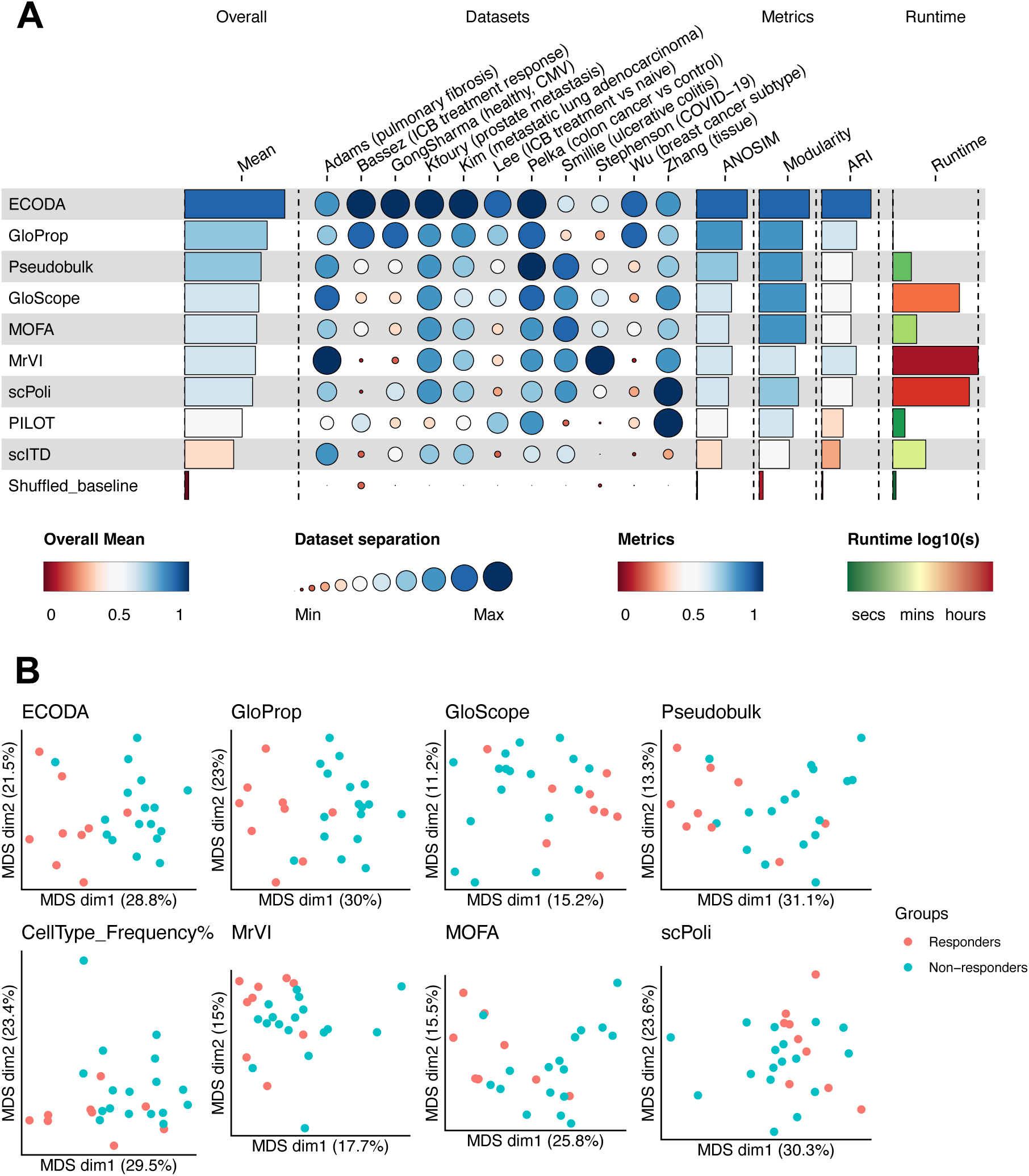
Benchmark of scRNA-seq sample representation methods for patient stratification. **A)** Representation methods are ranked by the average of three separation metrics: Analysis of Similarity (ANOSIM), graph Modularity and Adjusted Rand Index (ARI). The dots display average metric scores per dataset. Separation metrics were min-max-scaled across the tested methods per dataset to range from [0, 1], to give equal weights to each dataset. Unscaled metrics are shown in Supplementary figure 17. “Shuffled_baseline” is a negative control ECODA sample representation, where sample labels have been shuffled. **B)** Multidimensional Scaling (MDS) projections of a dataset of breast cancer patients (Bassez cohort) for a number of selected methods. Points represent individual patients (samples taken before treatment) and point color immunotherapy response status. Expanded benchmark results with additional methods and alternative ranking criteria are shown in Supplementary figure 15.

We also observed a substantial disparity in computational cost. While ECODA, GloProp and Pseudobulk embeddings were generated in seconds, GloScope, MrVI, and scPoli required hours of computation on GPU-enabled hardware; for these methods, it was necessary to downsample large cohorts to prevent out-of-memory errors (see Methods, Supplementary figure 14).

Notably, alternative cell-type proportion representations that do not use log-ratio-transformations, such as raw or arcsine-transformed cell-type frequencies, as implemented in previous studies (Yasumizu et al. 2024, Shitov, Dehkordi, and Luecken 2025, Phipson et al. 2022, Alayoubi, Bentsen, and Looso 2024) – performed substantially worse (Supplementary figure 15 “CellType_Frequency%”, Supplementary figure 16A “Frequency%”, and Supplemental Note). These results highlight the importance of properly handling cell-type proportions as compositional data (Greenacre 2021, Aitchison 1982). In contrast, performance was largely robust to the different zero-handling strategies used prior to log-ratio-transformation (Supplementary figure 16B and Supplemental Note).

The pseudobulk baseline representation, which captures differences in the average gene expression profile across all cells within a sample, showed a surprisingly high performance for unsupervised stratification. To disentangle the effects of transcriptional changes and compositional shifts, we calculated sample distances based on per-cell type pseudobulks (see Methods). This representation had markedly decreased performance compared to the global pseudobulk (Supplementary figure 15, “Pseudobulk_CT”), indicating that stratification performance is driven more by compositional differences than by within-cell-type transcriptional variability.

In summary, ECODA represents a top-performing, interpretable and scalable approach for unsupervised patient stratification.

### ECODA is robust to different cell type annotation strategies

A potential limitation of ECODA is its reliance on cell-type annotation. In our benchmark, ECODA was applied using author-provided cell-type labels (except for Lee and Zhang cohorts, for which such annotations were unavailable, see Methods). To evaluate the dependence of ECODA’s performance on expert-curated cell-type annotations, we compared stratification performance using alternative cell-type annotation strategies for all datasets. Specifically, we contrasted high-resolution author-provided cell type labels with i) unsupervised Leiden cell clustering at multiple resolutions and ii) automated annotation tools, including HiTME (Andreatta et al. 2021a, Andreatta, Berenstein, and Carmona 2022) and scATOMIC (Nofech-Mozes et al. 2023) (see Methods).

High-resolution author-provided cell-type labels (ECODA_authors_HR) yielded the highest average separation scores (Supplementary figure 15). However, labels derived from unsupervised Leiden clustering across multiple resolutions achieved comparable performance (Figure 3A, B, Supplementary figure 18). Cell-type labels obtained from automated annotation tools performed similarly or slightly worse on average than those derived from unsupervised clustering, likely because these tools were not trained on all relevant tissues represented in our benchmark. Nonetheless, automated annotation outperformed author-provided labels and unsupervised clustering in the Bassez, Lee and Stephenson cohorts (Supplementary figure 19). A significant drop in performance was observed when the annotation granularity was reduced to coarse cell-type groupings, including i) low-resolution author-provided labels (ECODA_authors_LR), ii) low-resolution Leiden clustering (ECODA_Leiden_res_0.1 resulting in about 10 cell types, similarly to ECODA_authors_LR), or iii) pseudobulk-based cell-type deconvolution (ECODA_deconv) (Figure 3A,B, see Methods).

**Figure 3:**
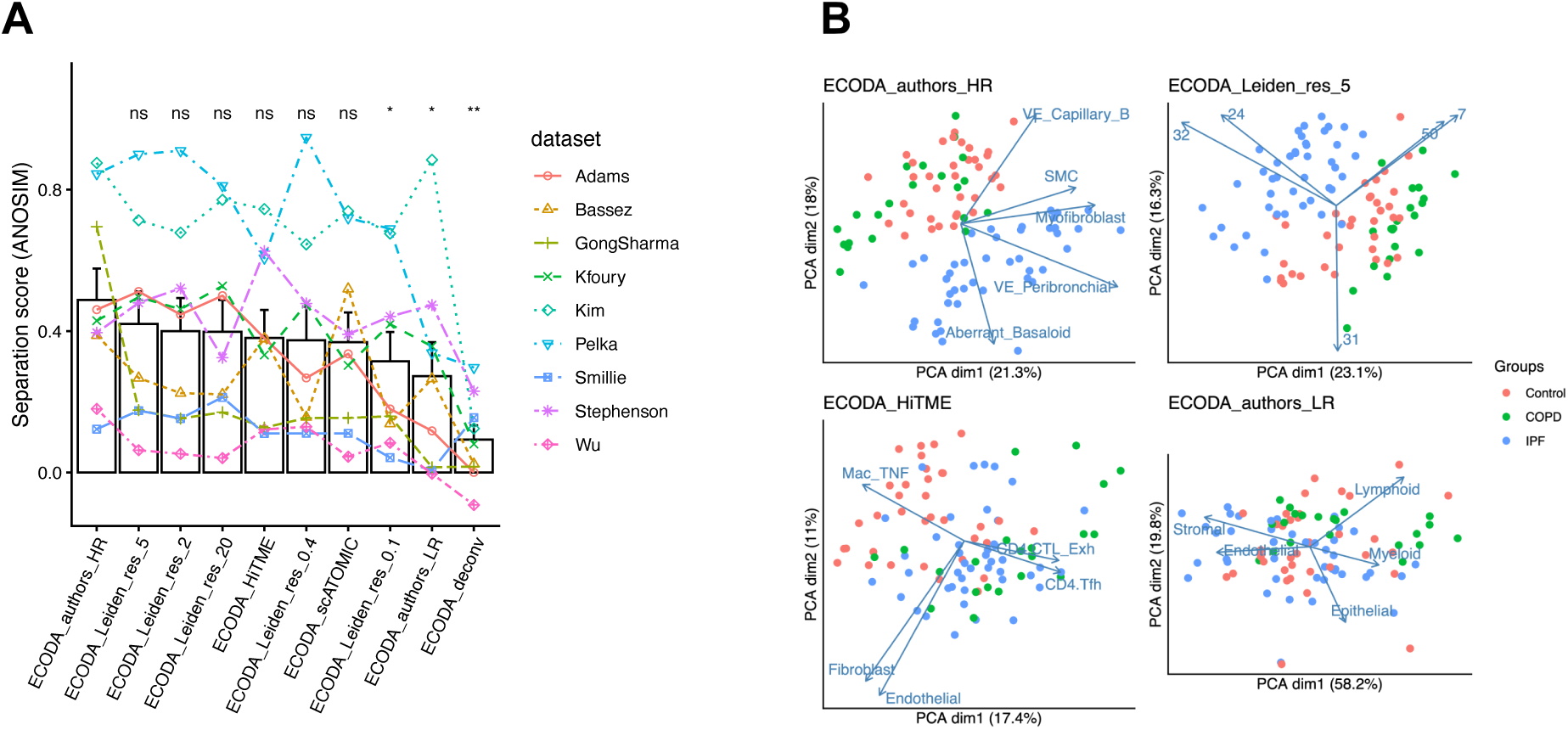
Impact of cell-type annotation granularity and method on ECODA performance. **A)** ANOSIM separation scores (y-axis) using ECODA based on different annotation strategies (x-axis) for the 9 datasets for which authors provided high-granularity cell labels. Annotation strategies include high-resolution (HR) and low-resolution (LR) author annotations, unsupervised Leiden clustering at various resolutions (0.1 to 20), and automated pipelines (HiTME, scATOMIC, deconvolution from pseudobulk). Asterisks indicate statistical significance from a paired Wilcoxon test against ECODA_authors_HR (ns: not significant, *: *p* < 0.05, **: *p* < 0.01). **B)** First principal components PC1 and PC2 for the Adams pulmonary fibrosis dataset based on four cell-type annotation methods: high-resolution author labels, unsupervised Leiden cell clustering, automated annotation using HiTME, low-resolution author labels. Arrows show top 5 features by explained variance for each annotation method (see Methods). COPD: chronic obstructive pulmonary disease, IPF: idiopathic pulmonary fibrosis, Control: healthy.

In summary, although ECODA relies on cell-type labels, it robustly captures cohort-level structure across a broad range of manual, unsupervised, and automated annotation strategies. This flexibility indicates that ECODA remains effective for patient stratification even in the absence of expert manual curation, provided that a sufficient level of cell-type granularity is available.

### ECODA performance is driven by a small subset of highly variable cell types

High-resolution cell-type annotation often consists of several dozen distinct populations. We investigated whether retaining only the most highly variable cell types (HVCs) is sufficient for patient stratification. Across datasets, we found that between 5 and 18 HVCs –corresponding to 12-29% of the total number of cell types per dataset– were sufficient to explain 40% of the total compositional variance (Figure 4A, Supplementary figure 20). In this range of variance explained, stratification performance remained stable (Figure 4B, Supplementary figure 21), indicating that ECODA’s performance is largely driven by a limited subset of highly variable cell types.

**Figure 4:**
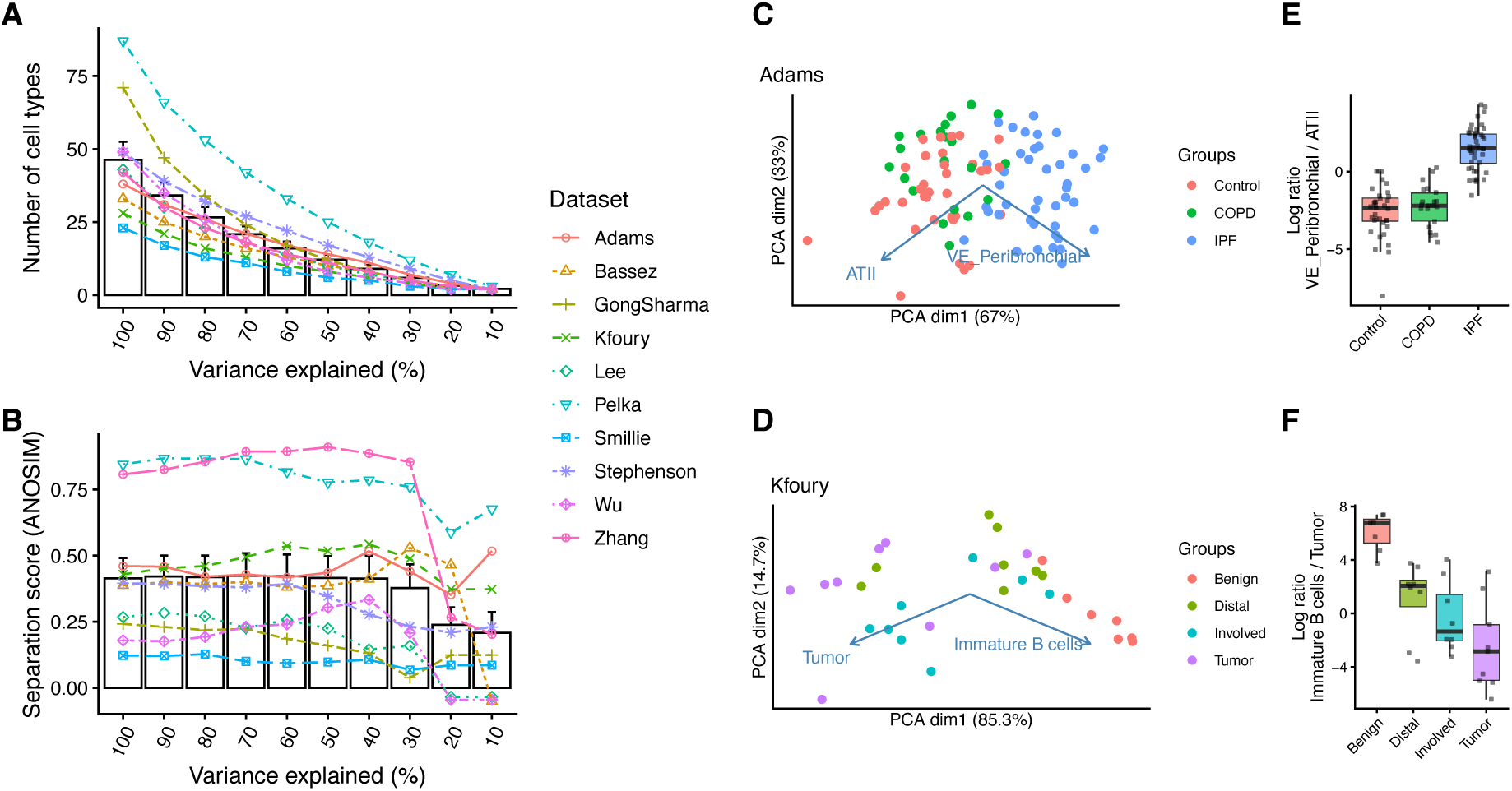
ECODA Performance is Driven by a Small Subset of Highly Variable Cell Types (HVC). **A)** The number of top HVCs (y-axis) required to reach specific cumulative variance explained (x-axis) on average across datasets. **B)** Separation performance (ANOSIM) as a function of retained variance. The bars represent the mean value. Whiskers show standard error of the mean. **C,D)** PCA biplots based on the CLR-transformed frequencies of the two most highly variable cell types for the Adams (ATII and VE_Peribronchial) and Kfoury (TIM and Immature B cells) datasets. **E,F)** Log-ratios of the two most highly variable cell types in the Adams and Kfoury datasets. COPD: chronic obstructive pulmonary disease, IPF: idiopathic pulmonary fibrosis, Control: healthy, Benign: healthy inflamed hip bone, Distal: liquid vertebral bone marrow distant from tumor, Involved: liquid vertebral bone marrow next to tumor, Tumor: solid metastatic tumor.

In some cases, a small number of cell types were sufficient to accurately separate biological conditions. For example, in the Adams dataset (pulmonary fibrosis), just two cell types – alveolar type 2 (ATII) and peribronchial vascular endothelial cells – were sufficient to separate samples by disease status (Normal, IPF, or COPD) (Figure 4C). In the Kfoury dataset (prostate cancer bone metastases), immature B cells and tumor inflammatory monocytes (TIMs) largely separated samples according to anatomical location and malignancy status (Figure 4D). In both cases, a simple score based on the cell counts ratio of two cell types – which could be quantified using low-plex staining platforms routinely employed in clinical practice – was sufficient to stratify patients according to relevant biological variables (Figure 4E,F).

These findings indicate that ECODA’s performance is primarily driven by signal concentrated in a small subset of biologically relevant cell populations. Moreover, they highlight the interpretability of compositional representations, which directly identify the specific cell types underlying cohort-level biological variation.

### ECODA is robust to batch effects

Technical batch effects can mask biological signals, particularly when confounded with experimental design. We hypothesized that cell-type compositions would be less sensitive to batch effects than gene expression profiles. To evaluate this, we examined a large cohort (Joanito et al. 2022) of samples from different types of tissue (colorectal tumors, adjacent normal tissue, and lymph nodes, Figure 5A,C), which were sequenced in two batches using different chemistries (3’ vs. 5’, Figure 5B,D).

**Figure 5:**
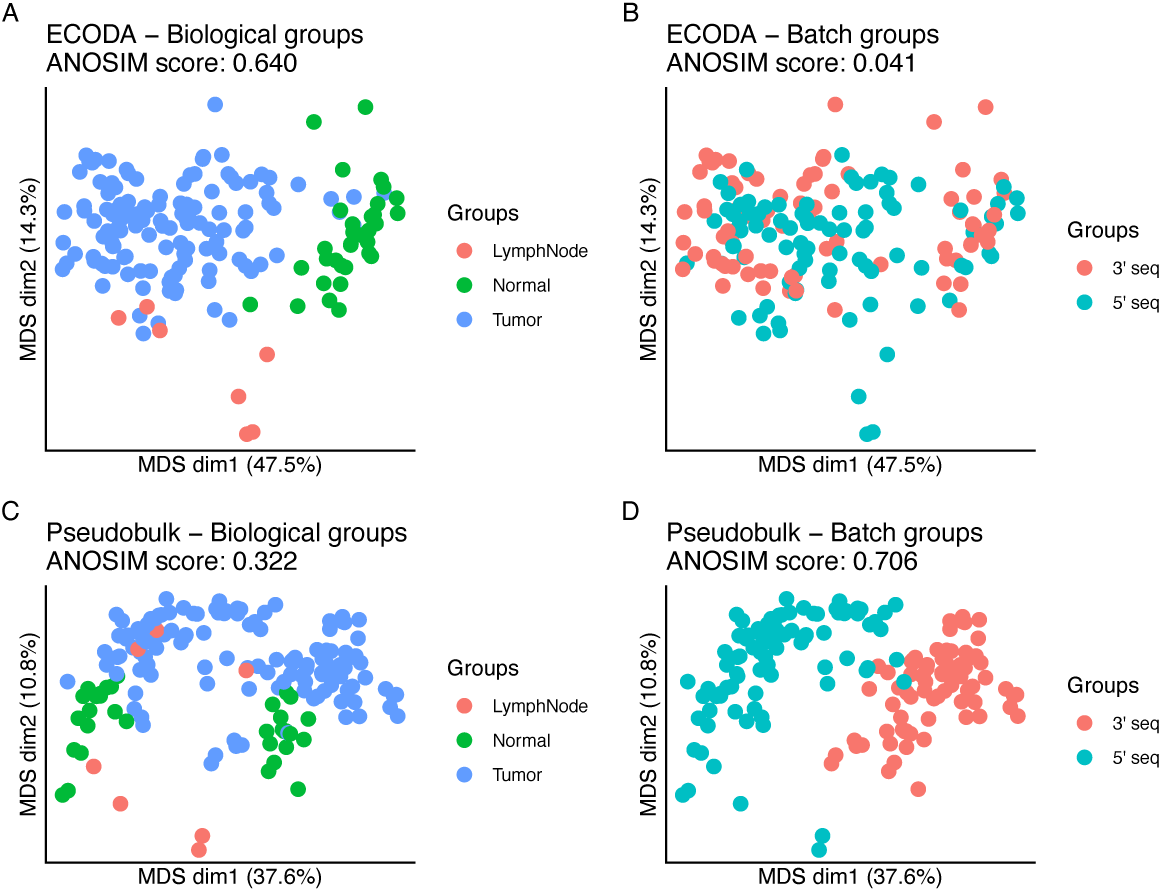
ECODA is robust to batch effects. Multidimensional Scaling (MDS) biplots of ECODA **(A, B)** and pseudobulk representation **(C, D)** showing sample separation by biological condition (Lymph Node, Normal, Tumor, **A, C**) and by technical batches (3’ vs. 5’ sequencing protocol, **B, D**) in the Joanito et al. dataset. Corresponding ANOSIM separation scores are shown. Results for another multi-batch cohort dataset are shown in Supplementary Figure 22.

In the ECODA representation, batch separation was modest (ANOSIM = 0.041, Figure 5B), whereas tissue-type separation was substantially stronger (ANOSIM = 0.640, Figure 5A). In contrast, the pseudobulk representation showed weaker biological (tissue-type) separation (ANOSIM = 0.322, Figure 5C) and pronounced batch separation (ANOSIM = 0.706, Figure 5D). Similar patterns were observed in an additional multi-study dataset (Supplementary figure 22 and Supplementary Table 4). These results indicate that cell type compositional embeddings are comparatively robust to batch effects while preserving biologically meaningful structure.

#### scECODA: a framework for single-cell exploratory cell-type compositional data analysis

To facilitate cohort-level exploratory analysis of scRNA-seq data, we developed scECODA (single-cell Exploratory COmpositional Data Analysis). scECODA transforms annotated single-cell gene expression objects from Bioconductor (Huber et al. 2015) or Seurat (Hao et al. 2024a) workflows into interpretable sample-level representations that capture biologically meaningful variation in cellular composition across samples.

For each sample, cell-type proportions are computed and transformed with the centered log-ratio (CLR). The resulting CLR-transformed cell-type proportions matrix constitutes the core sample-level representation, enabling distance calculations between samples in Euclidean space. From this representation, scECODA supports dimensionality reduction (e.g., principal component analysis), unsupervised clustering, and quantitative assessment of sample separation using metrics such as Adjusted Rand Index (ARI), graph Modularity, and ANOSIM. The package includes multiple visualization functions to explore cohort structure and associated biological variables. In addition to sample-level analysis, scECODA enables the investigation of cell-type co-variation patterns, revealing coordinated cellular responses and shared tissue microenvironments, and the identification of Highly Variable Cell types (HVCs), which capture the primary biological variance underlying sample stratification. Given the competitive performance of pseudobulk representations without requiring cell-type annotations, the scECODA package provides dedicated functionality for both compositional and pseudobulk-based cohort analyses.

Overall, scECODA provides a simple, scalable, and interpretable framework for exploratory cohort-level analysis and patient stratification based on single-cell transcriptomic data. scECODA is available as an R package at: https://github.com/carmonalab/scECODA.

## Discussion

As single-cell RNA sequencing technologies continue to improve in throughput, efficiency, and multiplexing capacity (e.g., 384-plex using the 10x Genomics FLEX assay), per-sample costs are decreasing, making large cohort-level profiling increasingly feasible. In this setting, exploratory analysis at the cohort level becomes essential to uncover latent biological structure, including patient subgroups with shared pathologies or treatment responses.

Here, we benchmarked current approaches for scRNA-seq-based patient stratification across 11 cohorts and covering a broad range of biological conditions. We observed a surprisingly strong performance of simple baseline methods: representations based on cell-type composition and whole-sample average (pseudobulk) gene expression matched or outperformed more complex state-of-the-art sample representation approaches. In particular, centered log-ratio (CLR)-transformed cell-type proportions – which account for the compositional constraint that proportions sum to one (here referred to as ECODA) – achieved the highest performance and increased robustness to batch effects. In contrast, cell-type proportion-based representations that do not use log-ratios – such as raw cell-type frequencies and GloProp, which compares samples as discrete cell-type distributions using a divergence on the proportion vectors – showed lower performances. Importantly, ECODA can produce sample embeddings even for large cohorts in a matter of seconds on a standard computer; competing methods require hours of computation and become unfeasible in terms of random-access memory (RAM) for cohorts of hundreds of samples.

The strong performance of ECODA suggests that, provided a set of fine-grained cell types, inter-sample biological variation is largely explained by shifts in cell-type abundance rather than by transcriptional reprogramming within those cell types. A second key insight is that sample separation is frequently driven by a limited subset of highly variable cell types. In many datasets, retaining approximately the top third of cell types by variance was sufficient to preserve discriminatory power, indicating that disease-associated variation may reflect specific cell type compositional signatures rather than widespread perturbations.

Beyond its superior performance, ECODA offers two key advantages: interpretability and translational relevance. Representations learned by deep neural networks are difficult to interpret biologically, whereas compositional representations directly identify the cell types responsible for inter-sample differences. For example, ratios between exhausted and naïve CD8+ T cells may provide actionable insight into immunotherapy response. Furthermore, compositional signals align naturally with established clinical platforms such as flow cytometry and immunohistochemistry. By identifying specific cellular drivers through ECODA, discovery-phase single-cell profiling can be translated into targeted, cost-effective diagnostic assays based on simple cell-type ratios, analogous to established biomarkers such as the neutrophil-to-lymphocyte ratio.

A potential limitation of ECODA is its reliance on cell-type annotations. However, our analyses demonstrate that performance is robust to the annotation strategy. Unsupervised Leiden clustering at sufficient resolution performed comparably to expert-curated labels, indicating that annotation granularity, rather than manual curation *per se*, is the primary determinant of biological separation. This suggests that ECODA can serve as an early exploratory step in unsupervised workflows, enabling identification of major axes of patient variation prior to detailed manual annotation. At the same time, in several datasets, expert-curated and automated supervised cell-type classification outperformed purely unsupervised clustering. This likely reflects the ability of biologically informed annotation strategies to resolve subtle but biologically meaningful distinctions that may be obscured in global transcriptional space. Thus, patient stratification performance using ECODA will benefit from improved methods for automated high-resolution cell-type classification.

In conclusion, we propose exploratory compositional data analysis as an accurate, scalable and highly interpretable approach for representing single-cell transcriptomics data at the cohort level. Because the principles of cell-type compositional analysis are modality-agnostic, ECODA can be extended to other technologies that quantify cellular composition, including flow cytometry, spatial omics, and high-plex imaging. In addition to enabling patient stratification, ECODA provides a practical quality-control framework to detect cohort structure and potential confounders, including batch effects, early in the analytical pipeline. To facilitate broad adoption, we provide scECODA as an open-source R package.

## Methods

### scRNA-seq data pre-processing

Count matrices of all datasets (see Supplementary table 2 for details on sample inclusion criteria) were log1p-normalized using the NormalizeData() function in Seurat (Hao et al. 2024b) v5 using default parameters. Cells were quality-filtered based on ribosomal and mitochondrial content and based on RNA counts distributions. After filtering, only samples with at least 500 cells were retained. Highly variable genes (HVGs) were identified using the Seurat function FindVariableFeatures() with default parameters. Principal component analysis (PCA) was performed on normalized, scaled expression of the top *N* HVGs (*N*=2000 unless specified in the text).

### Cell type annotation

To assess the impact of cell-type annotation methodology and granularity on sample representation, we compared three strategies:

- Expert author annotations: By default, we used the highest-granularity cell-type labels provided by the authors of the original studies. When available, we also compared different levels of resolution for cell-type annotation, e.g. “T cells” as low-resolution labels, and T cell subtypes for high-resolution labels. For the Kfoury dataset, high-resolution subtypes were manually consolidated into broader categories, as low-resolution labels were not provided by the authors. For the Lee dataset, author labels were not provided. For the Zhang dataset, cells from the two tissues (blood and tumor) had inconsistent cell-type labels and could not be compared. Thus, for the latter two datasets, automated annotation by HiTME was used (see below), and they were excluded from the analysis of the effect of cell-type annotation methods.
- Automated reference-based annotation: We used two automated annotation tools: i) HiTME (Cancer Systems Immunology Lab [2023] 2026) (a combination of scGate (Andreatta, Berenstein, and Carmona 2022) (human HiTME models) and ProjecTILs (Andreatta et al. 2021b, 2022) (sketched CD8T v1, CD4T v2, MoMac v1 and DC v2 maps)); and ii) scATOMIC (Nofech-Mozes et al. 2023) v2.0.3 with default parameters, using the option “normal_tissue” where applicable.
- Unsupervised cell clustering: Leiden clustering served as a biological knowledge-independent baseline. Clustering was performed within Seurat and default parameters on a nearest neighbor graph with the functions FindNeighbors() and FindClusters() using multiple alternative resolution values (0.1, 0.4, 2, 5, and 20).

### Patient stratification benchmark implementation

We benchmarked seven state-of-the-art methods for sample-level representation of scRNA-seq samples (MOFA+, scITD, GloScope, GloProp, MrVI, PILOT, scPoli), in addition to two baselines based on average gene expression profiles (Pseudobulk) or cell type composition (ECODA) (for specific method parameters used, see specific methods section below). Sample representation methods take as input a gene-by-cell gene expression matrix or a derived lower-dimensional embedding (scITD, GloScope, MrVI, PILOT, scPoli), a gene-by-sample average gene expression matrix (MOFA+, Pseudobulk), or a cell type-by-sample matrix (ECODA, GloProp). The methods produce either i) directly a sample-by-sample symmetric dissimilarity matrix encoding cohort-level structure, or ii) a reduced-dimensional sample embedding. An overview of the inputs and outputs for each method is provided in Supplementary table 3. For the methods that return sample embeddings, we calculated sample-by-sample (*N x N*) distance matrix by computing the pairwise Euclidean distance across all embedding dimensions using the stats::dist() function in R. This step ensures that all methods are represented in a unified distance-matrix format, allowing for a direct comparison using the same downstream evaluation metrics. Based on the distance/dissimilarity matrix obtained for each method, separation of biologically relevant groups of samples was quantified using ANOSIM, Modularity and ARI (see Methods section on “Separation quantification metrics”). To visualize the sample-level relationships, we applied Classical Multidimensional Scaling (MDS), also known as Principal Coordinates Analysis (PCoA), using the stats::cmdscale() function in R. Unlike PCA, which requires a starting feature matrix, MDS operates directly on the calculated dissimilarity matrices, allowing to visualize all methods. It projects the *N x N* distances into a 2D coordinate system while preserving the global pairwise distances as accurately as possible. The axes were scaled by the eigenvalues of the decomposition to represent the percentage of variance explained. To accommodate memory-intensive methods, we downsampled the Gong & Sharma dataset to a maximum of 5000 cells per sample. We further controlled this specific dataset for demographic confounding by restricting the cohort to 180 males aged ≤ 40 years. This refinement isolates the immunological signature of cytomegalovirus (CMV) infection, ensuring that observed variance stems from CMV status rather than age or sex.

Execution times were measured on a desktop computer (16-core AMD Ryzen 9 5950X, 128 GB RAM and additional 300 GB swap memory on Samsung 990 PRO 2TB SSD, NVIDIA GeForce RTX 4090 GPU). GPU-capable methods were executed on the GPU, with the exception of MrVI, which was run on the CPU due to persistent stability issues during GPU execution and MOFA+ which was reasonably fast already on the CPU for the data utilized in this study. Execution times exclude standard upstream processing (e.g., PCA and cell type annotation) to focus on the computational overhead specific to each method. Consequently, only tasks required for downstream integration, such as pseudobulk generation or sample-level embedding, were included in the final timing analysis.

### Sample representation methods configuration

Gene expression-based methods (Pseudobulk, MOFA+, scITD, GloScope, MrVI, PILOT, scPoli) were run on the 2000 most highly variable genes (HVG). For Pseudobulk and MOFA+, HVGs were identified from the aggregated pseudobulk matrix using the dispersion-based ranking provided by DESeq2 (Love, Huber, and Anders 2014). For the other methods working directly on the gene-by-cell gene expression matrix (scITD, GloScope, MrVI, PILOT, scPoli) Seurat’s FindVariableFeatures() function was used (see Method section “scRNA-seq data pre-processing”).

ECODA: Cell counts for each cell type were transformed using Centered Log-Ratio (CLR) transformation *clr*(*x*) (see Supplemental Note) by dividing each component of a vector *x* by the geometric mean of all components in the vector *g*(*x*)and taking the natural logarithm:

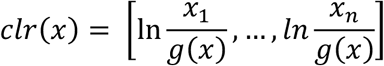

To avoid zeros, a pseudocount of 0.5 was added to zero-count values (Martín-Fernández, Barceló-Vidal, and Pawlowsky-Glahn 2003, Greenacre 2021, Lubbe, Filzmoser, and Templ 2021) (used for benchmark). Other zero handling methods were tested (see Supplemental Note).

Whole-sample pseudobulk: Raw counts were summed across all cells per sample and normalized using DESeq2 v1.44.0 (Love, Huber, and Anders 2014). We utilized the median ratio method for size factor estimation followed by variance stabilizing transformation.

Deconvoluted Cell type Composition from pseudobulk: Raw pseudobulk counts were deconvoluted with EPIC v1.1.7 using the blood immune reference profiles (BRef) (Racle and Gfeller 2020). Zeros in percentages were replaced (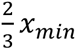, where *x_min_* is the lowest cell type abundance value (in frequency%) in the dataset) and CLR-transformed.

All state-of-the-art methods were run with author-suggested defaults. Additionally, to evaluate the robustness and ensure a fair comparison, all tested methods were run across a range of their respective most important parameters. For example, for gene expression-based methods, we varied the number of HVGs (1000, 2000, 3000) (for benchmark: 2000 HVGs) and number of extracted embedding dimensions. Additional important parameters for each method are listed below:

- ECODA: The CLR-transformed counts (sample x cell type abundance matrix) (for benchmark) or the first 2, 3, 5, 10, or 15 PCA components calculated from cell type-wise centered unscaled CLR-transformed abundances (sample x PCA component matrix) were used to calculate the pairwise Euclidian sample distance matrix.
- Pseudobulk: Pairwise Euclidian sample distances were computed either directly from the DESeq2-pre-processed pseubodulk HVG expression matrix (for benchmark), or the first 2, 3, 5, 10, 15, 30 or 50 PCA components calculated from the gene-wise centered unscaled DESeq2-pre-processed pseubodulk HVG expression matrix.
- Pseudobulk per cell type (“pseudobulk_CT”): This approach aims to capture transcriptional differences between samples that are independent of cell type composition. For each cell type, a DESeq2-preprocessed gene-by-sample pseudobulk expression matrix was generated, including only samples in which the cell type was detected (with a minimum of 5 cell counts). Each matrix was subset to the top 500 or 2000 highly variable genes (HVGs), and a sample-by-sample Euclidean distance matrix (*D_c_*) was computed, for each cell type. The combined distance matrix 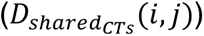 was then obtained by averaging distances across shared cell types. Specifically, for any pair of samples *i* and *j*, the distance was defined as the sum of their distances across all shared cell types divided by the number of shared cell types.

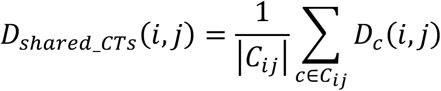

where *C_ij_* is the set of cell types shared between samples *i* and *j*. |*C_ij_*| is the number of shared cell types. *D_c_*(*i*, *j*) is the Euclidean distance between samples *i* and *j* computed from the pseudobulk matrix of cell type *c*.

- MOFA+: Run on DESeq2-normalized pseudobulk HVGs; 2, 3, 5, 10, 15 factors extracted from the “Z” matrix (for benchmark: 15 factors). The embedding matrix was used to calculate pairwise Euclidian sample distances.
- scITD: Coarse-grained author cell type labels were used; cell types with <5 counts in >20% of samples were omitted; 2, 3, 5, 10, 15 factors extracted from the sample scores matrix can in pbmc_container[[“tucker_results”]][[1]] (for benchmark: 5 factors). Coarse-grained cell type labels were used to limit sample dropout, as scITD drops samples that do not have at least 5 cells in each cell type; for the same reason, cell types with <5 counts in >20% of samples were omitted; 2, 3, 5, 10, 15 factors extracted from the sample scores matrix can in pbmc_container[[“tucker_results”]][[1]] (for benchmark: 5 factors). The embedding matrix was used to calculate pairwise Euclidian sample distances.
- GloScope: GloScope was run using the top 10, 30 or 50 PCA dimensions (for benchmark: 30 PCA dimensions); density estimation via k-Nearest Neighbors (k=25); distance via Kullback-Leibler (KL) divergence; distance matrix as square root of KL divergence. NA values were replaced by zeros as recommended by the authors.
- GloProp: Fine-grained author annotations were used in the function gloscopeProp (with ep = 0.5 and dist_metric = Kullback-Leibler (KL)). The final distance matrix was calculated as square root of the KL divergence matrix.
- MrVI: Trained for 50 epochs (convergence confirmed after 20 epochs). The resulting MrVI sample distance matrix was used directly.
- PILOT: 50 PCA components and fine-grained author cell type labels were used for the benchmark. Coarse-grained cell type labels were tested for the parameter screening but performed worse (Supplementary figure 2). The resulting PILOT sample Wasserstein distance matrix was used directly.
- scPoli: Embedding dimensions screened across 2, 3, 5, 10, 15 (for benchmark: 15 embedding dimensions). Fine-grained author cell type labels were used. Coarse-grained cell type labels were tested for the parameter screening and performed similarly (Supplementary figure 2). The embedding matrix was used to calculate pairwise Euclidian sample distances.

When the sample-by-PCA-loadings matrix was used (for ECODA or Pseudobulk), the distance matrix was calculated from the scores matrix calculated with the *prcomp* function from the stats R package.

Separation quantification metrics

To robustly quantify the separation of biological groups across different representations, we employed three complementary metrics: Analysis of Similarities (ANOSIM), Modularity, and the Adjusted Rand Index (ARI).

- ANOSIM: A rank-based metric that is parameter-free and robust to outliers. We utilized the anosim function from the vegan (Oksanen et al. 2025) R package, as established in previous benchmarks (Wang et al. 2024).
- Modularity (*Q*): A graph-based metric calculated using three nearest neighbors. As the maximum modularity (*Q*_%*3_) depends on the number of groups (Fortunato and Barthélemy 2007), we adjusted the modularity score (*Q_max_*) by the number of groups (*m*):

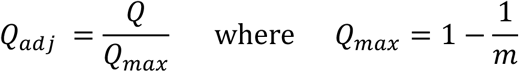

to ensure comparability across datasets with varying numbers of biological conditions.

- Adjusted Rand Index (ARI): A clustering-based metric (Hubert and Arabie 1985) that reflects real-world unsupervised exploratory workflows. ARI was calculated following hierarchical (Ward.D2, using the R package “stats” cutree function) or k-medoids clustering (using the R package “cluster” pam function), providing as input the known number of clusters, i.e. the actual number of ground-truth biological categories.

The metrics strongly correlated with the global average (mean across the three metrics) and with each other (Supplementary figure 1). ARI showed the highest correlation with the global mean (*R*^2^ = 0.90) followed by ANOSIM (*R*^2^ = 0.89). ANOSIM and ARI showed a consistent separation scoring (*R*^2^ = 0.81).

For the main benchmark, each metric score was min-max scaled to range between 0 and 1 per dataset. This normalization ensures that each dataset contributes equally to the combined benchmark score. Additionally, individual metric scores were aggregated across datasets by taking i) the average of the three min-max-scaled metric scores (Figure 2 and Supplementary figure 15), or ii) the average of the rank based on metric score (Supplementary figure 15).

### Selection of highly variable cell types

To focus on the most informative biological signals, we identified Highly Variable Cell types (HVCs) through a variance-based thresholding approach (for an example, see Supplementary figure 20). Raw counts were CLR-transformed after addition of a pseudocount, and the variance for each cell type across all samples was calculated and ranked in descending order. We selected the minimum number of top-ranked cell types required to reach a predefined cumulative variance threshold (e.g., 80% of total variance). To ensure a valid compositional representation, a minimum of two cell types was required for any dataset. Once HVCs were identified, the CLR-transformation was recalculated specifically on this subset.

### PCA and biplots: Identification of top contributing variables

To simultaneously display observations (sample embeddings) and variables (cell types or genes), we generated “biplots” using the factoextra R package (Kassambara et al. 2026). To identify the *n* most highly contributing variables for visualization, we used the select.var parameter in the fviz_pca function. Variables were ranked by their contribution to the principal components, which is defined as the ratio of their squared correlation with the components to the corresponding eigenvalues (Abdi and Williams 2010). This metric ensures that the selected features are those that explain the greatest proportion of variance in the displayed dimensions.

## Data availability

R-format RDS files for each dataset containing the respective Seurat object, including UMI count matrices, dataset-specific metadata, and cell type labels, are available at https://doi.org/10.6084/m9.figshare.31456522.

## Code availability

The code to reproduce all results in this study – including dataset pre-processing, reproducing the figures and the benchmark – is available at: https://github.com/carmonalab/ECODA_paper

The R package scECODA to perform cohort-level single-cell exploratory compositional data analysis for scRNA-seq data based on CLR-normalized cell type abundances is available at: https://github.com/carmonalab/scECODA

## Acknowledgements

This work was supported by the Swiss Cancer Research Foundation grant KFS-5409-08-2021 (S.J.C)

## Supplementary material

## Supplemental Note

### Normalization for cell-type compositional data

To identify the most effective way to represent cell-type composition, we compared several normalization and transformation strategies, including raw counts, frequencies, and log-ratio transformations. We found that the choice of normalization has a substantial impact on the resulting sample grouping structure (Supplementary figure 16A). Raw cell counts performed poorly, as expected from the large technical variation of the total cell numbers across samples. While converting counts to frequencies improved biological separation, log-ratio transformations significantly outperformed these naïve approaches, consistent with the constant-sum constraint in compositional data (Quinn et al. 2019, Greenacre et al. 2023). Compositional data needs to be brought from simplex to Euclidean space for distance calculations, which can be achieved by log-ratio transformation methods (Filzmoser, Hron, and Templ 2018, Lubbe, Filzmoser, and Templ 2021). Specifically, Centered Log-Ratio (CLR) transformation provided the highest mean separation score and was significantly more effective than arcsine square-root (*p* < 0.01, Wilcoxon paired test), additive log-ratio (ALR) with a low-variance reference (ALR_minCVref), or frequencies (*p* < 0.001, Wilcoxon paired test). ALR with a random cell type as reference (ALR_randref) did not perform worse (average of 20 random reference selections). However, CLR proved more practical than Additive Log-Ratio (ALR), as it avoids the requirement of selecting a specific reference cell type while maintaining comparable or superior performance.

Zeros are common in single-cell omics-derived cell-type compositions, since the typical number of cells profiled in each sample (∼10^3^) is not enough to detect low-frequency populations. Zeros prevent log calculation in the log-ratio transformations. Thus, we evaluated several zero-handling methods (Lubbe, Filzmoser, and Templ 2021), including simple pseudocount additions (Martín-Fernández, Barceló-Vidal, and Pawlowsky-Glahn 2003) (e.g. adding a count of one to all zeros or to all parts) and more complex model-based approaches (Palarea-Albaladejo and Martín-Fernández 2015, Templ, Hron, and Filzmoser 2011) like Multiplicative Log-normal replacement (multLN) and Multiplicative Replacement (multRepl) at varying thresholds (0.01% to 1%). In contrast to the primary normalization method, we found that the choice of zero-handling strategy had a relatively low impact on final separation scores (Supplementary figure 16B). Simple pseudocount methods such as replacing zeros with 1 (counts_zeros_1) or with 2/3 of the smallest value (i.e. 0.67 cell count), or adding 1 to all values (counts_all_1) performed highest and similarly to more complex model-based approaches, though a significant decrease in performance was observed when using multiplicative replacement at higher thresholds of 0.1% and 1% (*p* < 0.05, Wilcoxon paired test).

These results suggest that while log-ratio transformation is critical for unlocking the biological signal in compositional data, a simple pseudocount addition is a useful approach for avoiding zeros.

---

**Supplementary table 1:**
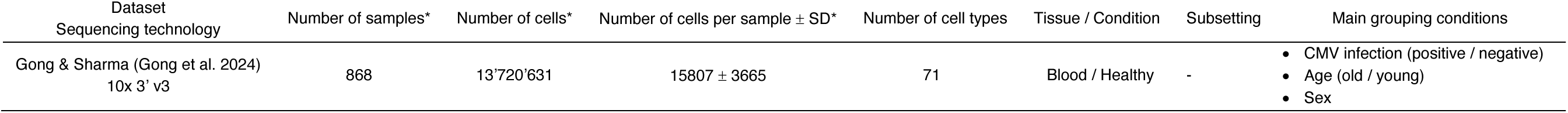
Gong & Sharma dataset used for Figure 1B. *: After quality filtering and subsetting. SD: standard deviation.

| Dataset<br>Sequencing technology | Number of samples* | Number of cells* | Number of cells per sample ± SD* | Number of cell types | Tissue / Condition | Subsetting | Main grouping conditions |
| --- | --- | --- | --- | --- | --- | --- | --- |
| Gong & Sharma (Gong et al. 2024)<br>10x 3' v3 | 868 | 13'720'631 | 15807 ± 3665 | 71 | Blood / Healthy | - | • CMV infection (positive / negative)<br>• Age (old / young)<br>• Sex |

**Supplementary table 2:**
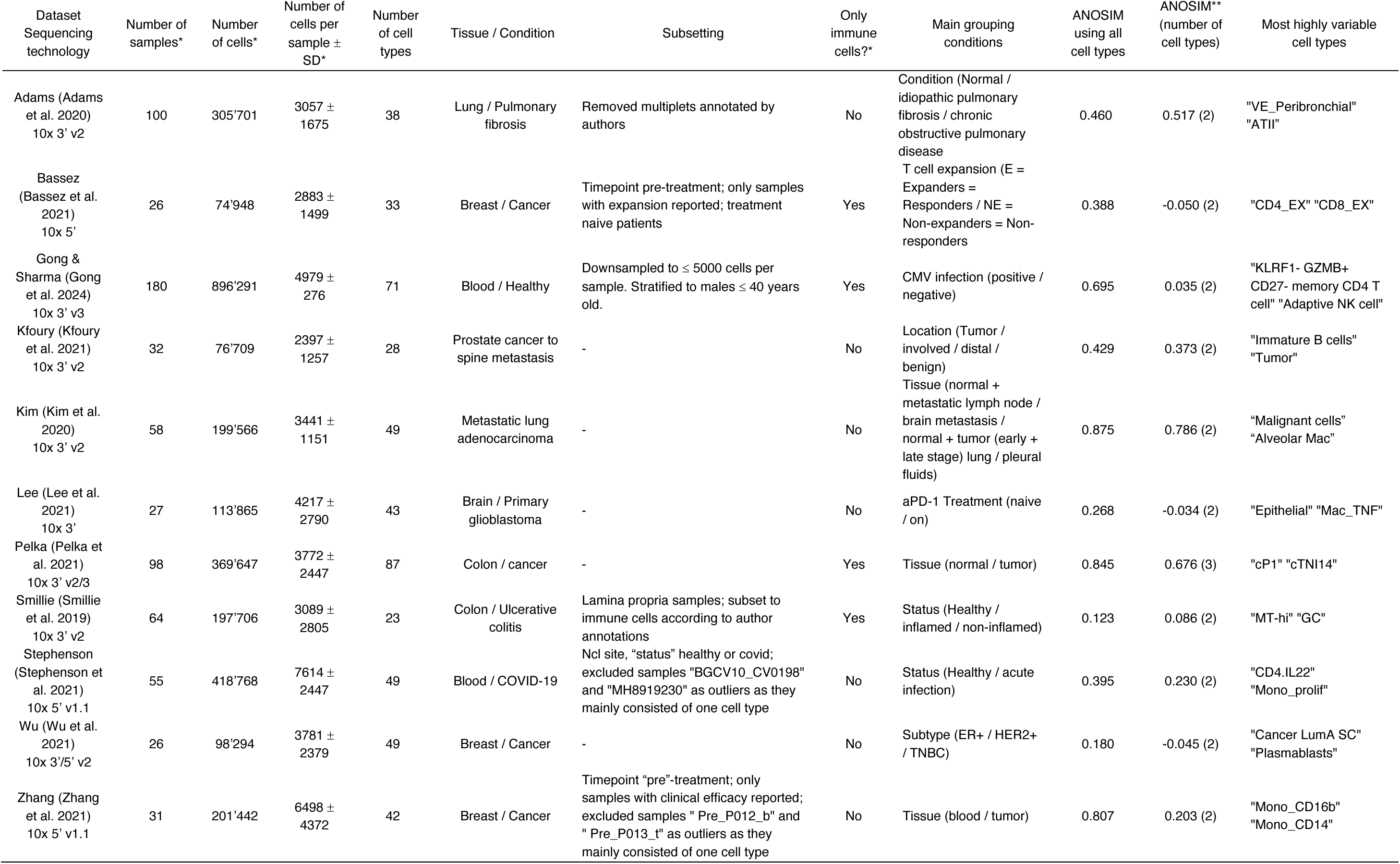
Datasets used for the sample embedding method benchmark. *After quality filtering and subsetting. SD: standard deviation. ** at 10% variance explained.

**Supplementary table 3:** Overview of available tools to create low-dimensional sample representations for exploratory analysis of scRNA-seq data. “Cell types” are defined by (e.g. automated or manual) annotation, giving each cell a type label. n = number of samples. HVGs: highly variable genes. “Factors” are extracted gene programs defined by the corresponding methods (scITD and MOFA+).

| Family | Name | Description | Input | Requires cell type annotation | Output |
| --- | --- | --- | --- | --- | --- |
| Cell type composition | ECODA | CLR-transformed cell type proportions | Cell type counts | Yes | Sample embeddings<br>($n \times \text{cell types}$) |
| | GloProp<br>(Wang et al. 2024) | Pairwise sample divergences, based on gene expression | Cell type counts | Yes | Distance matrix<br>($n \times n$) |
| Gene expression | Pseudobulk | Sample-aggregated counts | (DESeq2-pre-processed) sample bulk count matrix (2000 HVGs) | No | Sample embeddings<br>($n \times \text{HVGs}$) |
| Factor decomposition | scITD<br>(Mitchel et al. 2022) | Tensor factorization | Single-cell count matrix (here 2000 HVGs) and cell type annotation | Yes | Sample embeddings<br>($n \times \text{factors}$) |
| | MOFA+<br>(Argelaguet et al. 2018, 2020) | Matrix factorization | (DESeq2-pre-processed) bulk count matrix (here 2000 HVGs) | No | Sample embeddings<br>($n \times \text{factors}$) |
| Deep generative model | MrVI<br>(Boyeau et al. 2024) | Variational autoencoder | single-cell count matrix (here 2000 HVGs) | No | Distance matrix<br>($n \times n$) |
| | scPoli<br>(De Donno et al. 2023) | Variational autoencoder | single-cell count matrix (here 2000 HVGs) and cell type annotation | Yes | Distance matrix<br>($n \times n$) |
| Distributional representation | GloScope<br>(Wang et al. 2024) | Pairwise sample divergences, based on gene expression | PCA sample embeddings (here top 30 PCAs) | No | Distance matrix<br>($n \times n$) |
| | PILOT<br>(JoodakiMehdi et al. 2023) | Optimal transport / Variational Autoencoder | PCA sample embeddings (here top 50 PCAs) and cell type annotation | Yes | Distance matrix<br>($n \times n$) |

**Supplementary table 4:** Datasets used for batch effect testing. *: After quality filtering and subsetting. SD: standard deviation. *^$^*: Cell types obtained from annotation with HiTME that were present across all three datasets.

| Dataset<br>Sequencing technology | Number of<br>samples* | Number of<br>cells* | Number of cells<br>per sample ± SD* | Number of cell<br>types | Tissue / Condition | Subsetting | Main grouping<br>conditions |
| --- | --- | --- | --- | --- | --- | --- | --- |
| Joanito (Joanito et al. 2022)<br>10x 3' v2 and v3 / 10x 5' | 168 | 367'430 | 2187 ± 1295 | 11 | Lymph node / Normal /<br>Tumor / 3' seq / 5' seq | - | Tissue / Sequencing<br>technology |
| Gong & Sharma (Gong et al.<br>2024) 10x 3' v3 | 15 | 75'000 | 5000 ± 0 | 32 (HiTME<br>annotations) | Blood / Healthy | Downsampled to ≤ 5000 cells per sample. Stratified to males ≤ 40 years<br>old. Subset to the first 15 randomly selected samples. | Dataset |
| Stephenson (Stephenson et<br>al. 2021) 10x 5' v1.1 | 55 | 418'768 | 7614 ± 2447 | 32 (HiTME<br>annotations) | Blood / COVID-19 vs<br>healthy | Nci site, "status" healthy or covid; excluded samples "BGCV10_CV0198"<br>and "MH8919230" as outliers as they mainly consisted of one cell type | Dataset |
| Zhu (Zhu et al. 2023) 10x 3'<br>v2 | 17 | 129'939 | 7643 ± 1116 | 32 (HiTME<br>annotations) | Blood / Healthy | - | Dataset |

**Supplementary Figure 1:**
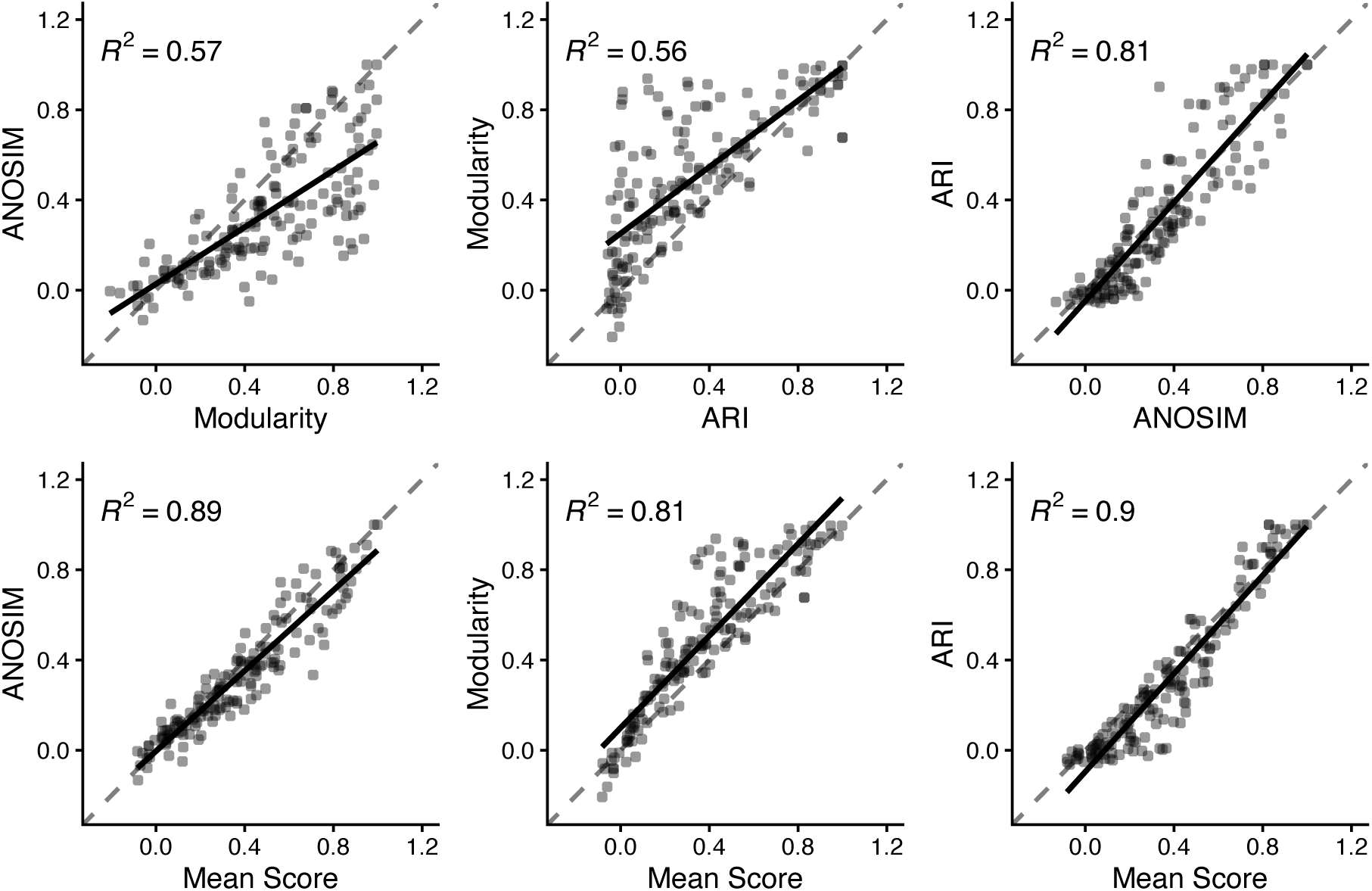
Metric Consistency Analysis. Scatter plots displaying the relationship between individual separation metrics (top) and the aggregate mean score (bottom) across all benchmarked methods and datasets. Each dot is the separation score for one benchmarked method (from Supplementary figure 15) on one dataset (Supplementary table 2). The solid black line represents the linear regression line, from which the coefficient of determination (*R^2^*) for each graph was derived. The dashed line represents equality line.

**Supplementary Figure 2:**
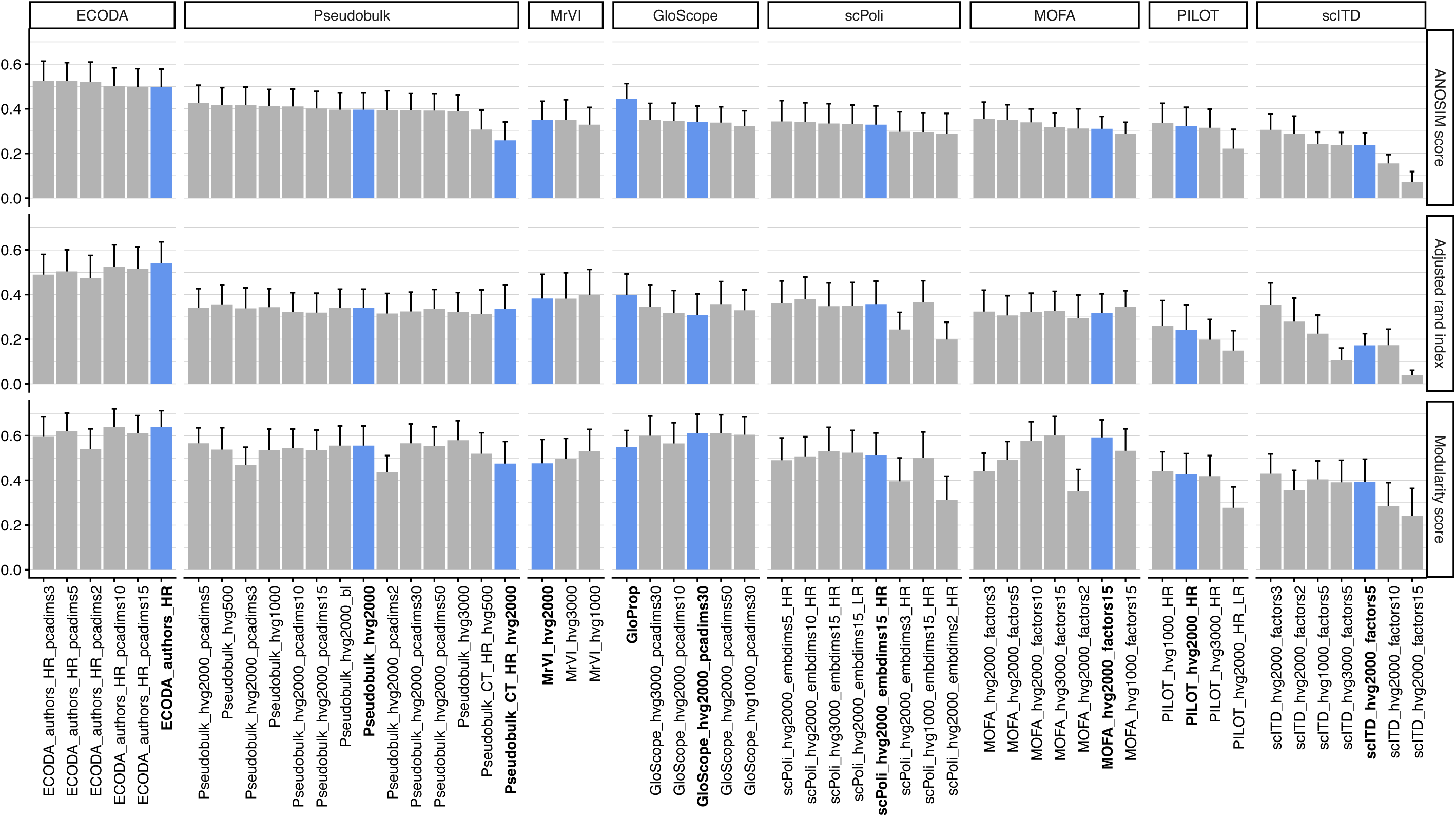
Parameter screening for benchmarked methods. To evaluate the robustness and ensure a fair comparison, all tested methods were run across a range of the most important parameters. Resulting scores for every method parameter combination tested are grouped by method and sorted by ascending ANOSIM score. Bars represent the mean separation score (y-axis) of each method with the indicated parameter setting, run across all benchmark datasets (Supplementary table 2). Whiskers represent standard error of the mean. Blue bars and bold text indicate that the configuration is displayed in the benchmark and extended benchmark figures.

**Supplementary Figure 3:**
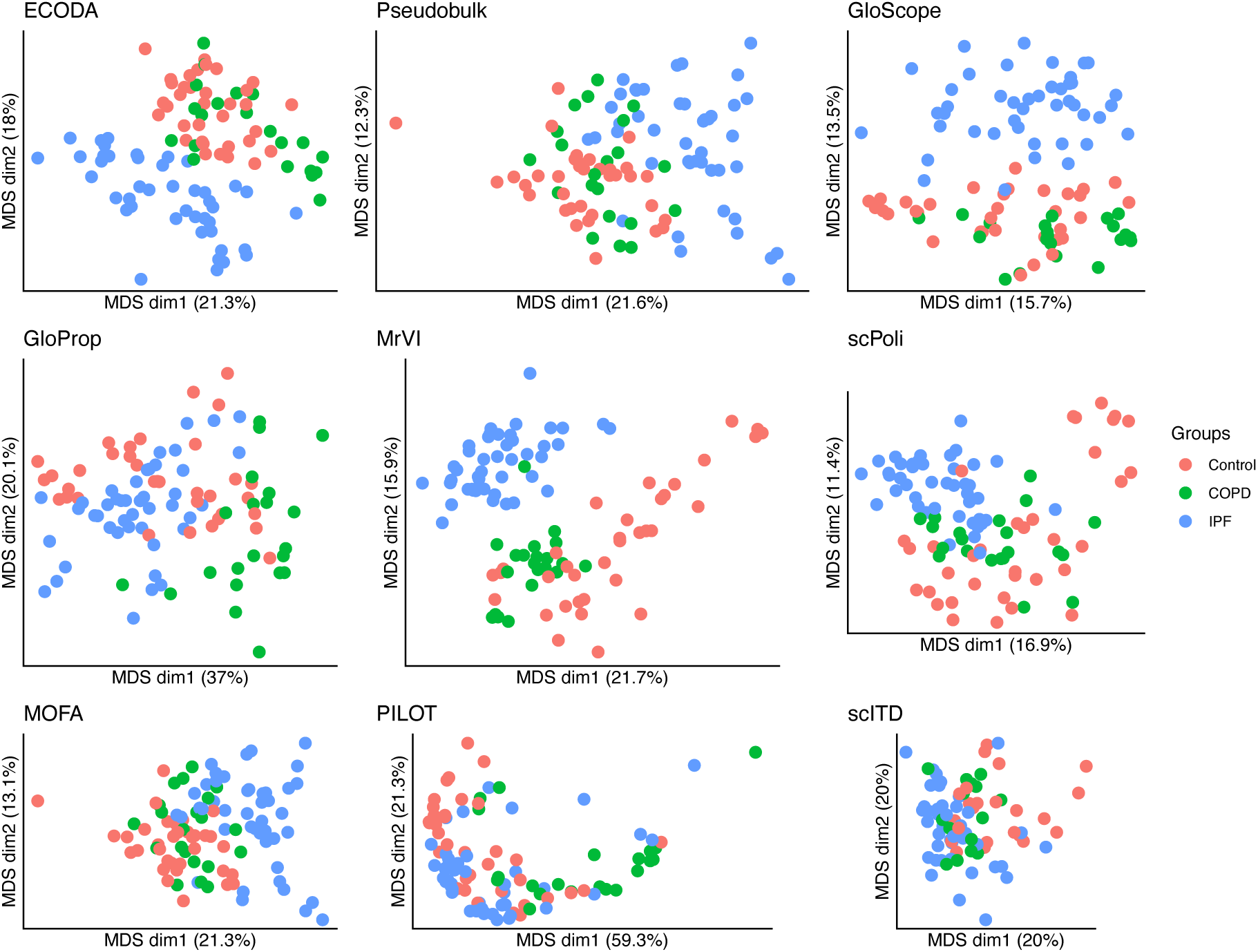
Classical Multidimensional Scaling (MDS) plots of inter-sample distance matrix of the Adams dataset. The groups in the Adams pulmonary fibrosis dataset are healthy control, chronic obstructive pulmonary disease (COPD) and idiopathic pulmonary fibrosis (IPF) samples.

**Supplementary Figure 4:**
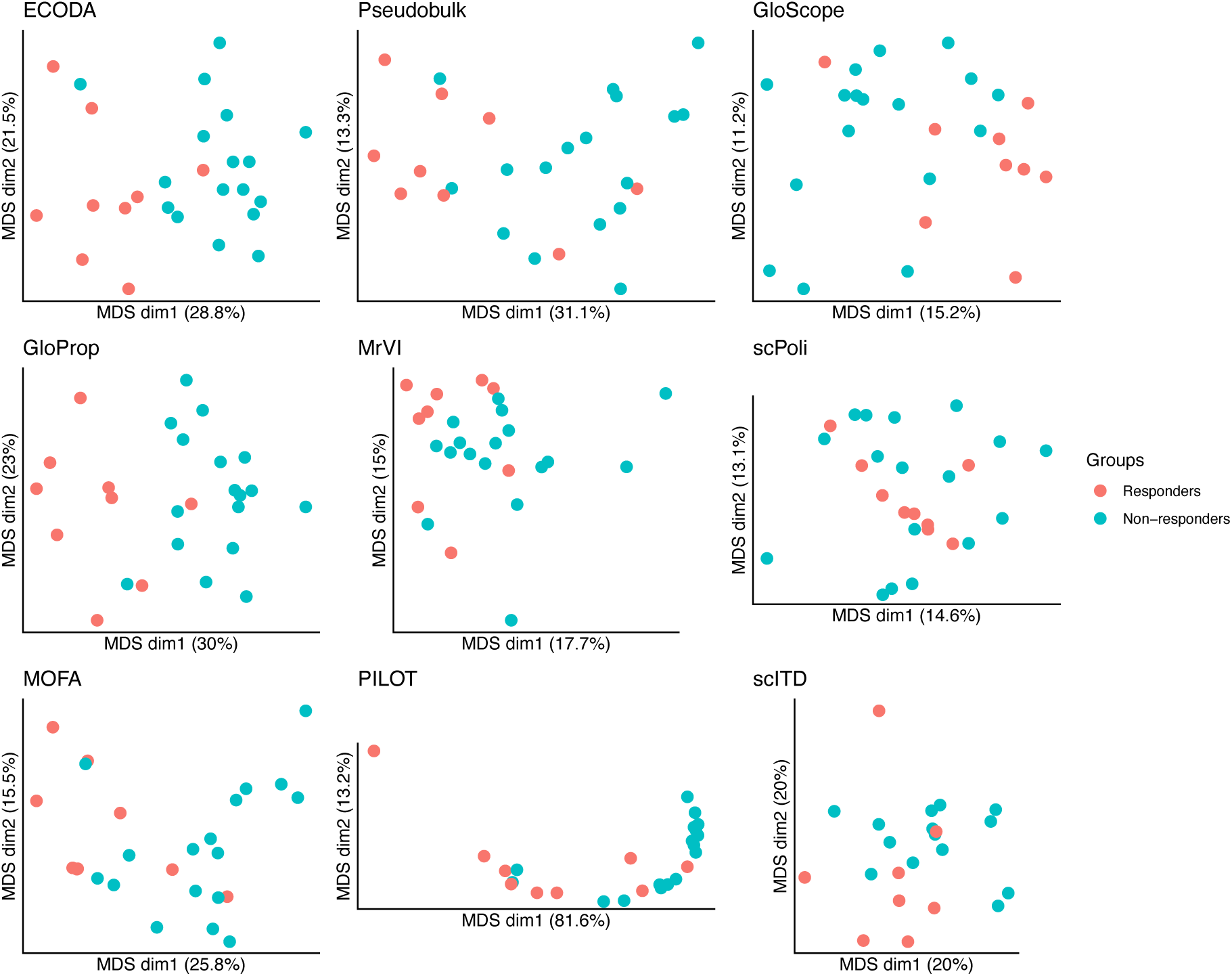
Classical Multidimensional Scaling (MDS) plots of inter-sample distance matrix of the Bassez dataset. The groups in the Bassez breast cancer dataset are responders and non-responders to immune checkpoint blockade treatment, respectively.

**Supplementary Figure 5:**
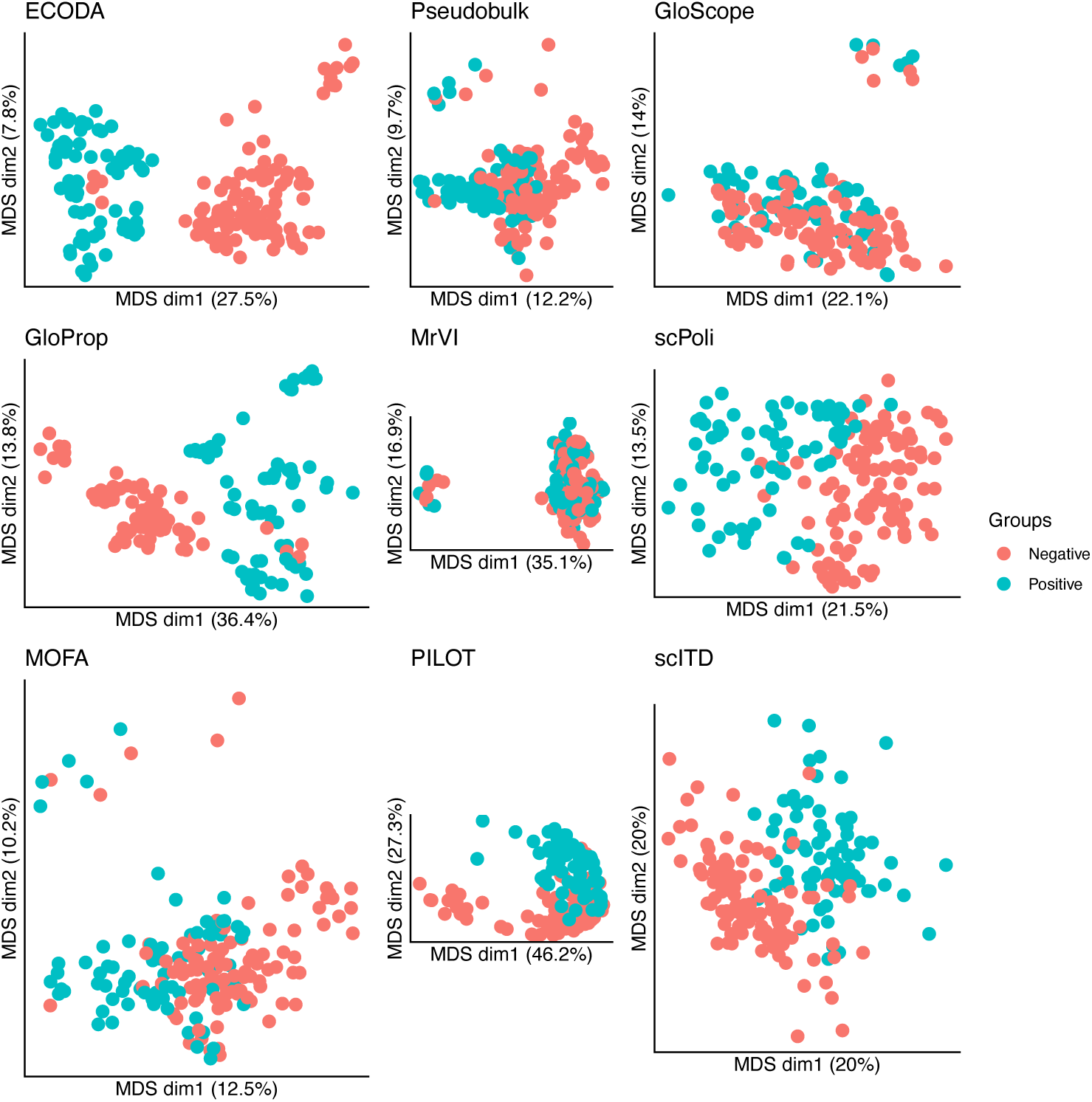
Classical Multidimensional Scaling (MDS) plots of inter-sample distance matrix of the Gong & Sharma dataset. The healthy cohort by Gong & Sharma, grouped by cytomegalovirus serology status.

**Supplementary Figure 6:**
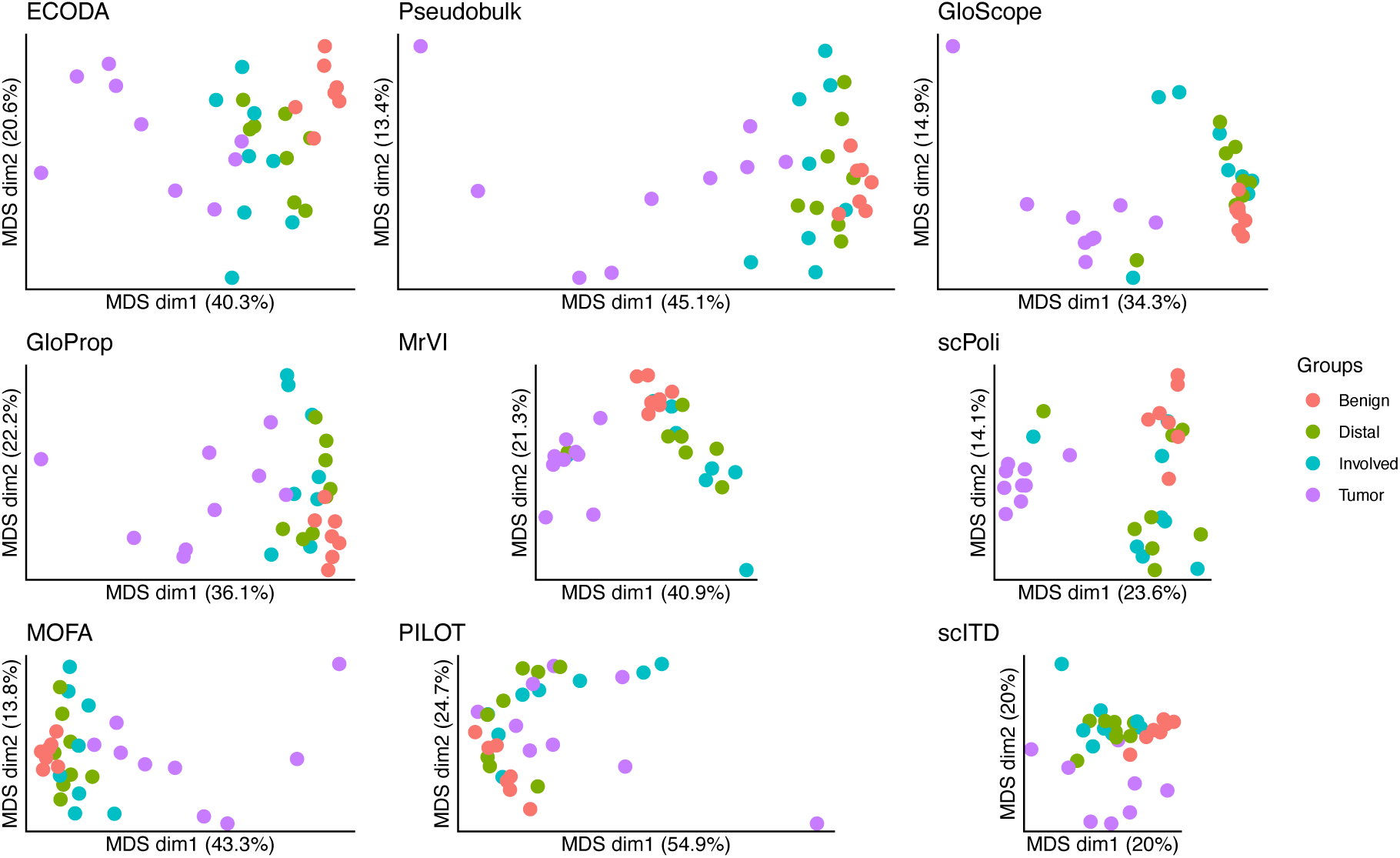
Classical Multidimensional Scaling (MDS) plots of inter-sample distance matrix of the Kfoury dataset. The groups in the Kfoury prostate cancer to spine metastasis dataset are from different locations: Benign: healthy inflamed hip bone, Distal: liquid vertebral bone marrow distant from tumor, Involved: liquid vertebral bone marrow next to tumor, Tumor: solid metastatic tumor.

**Supplementary Figure 7:**
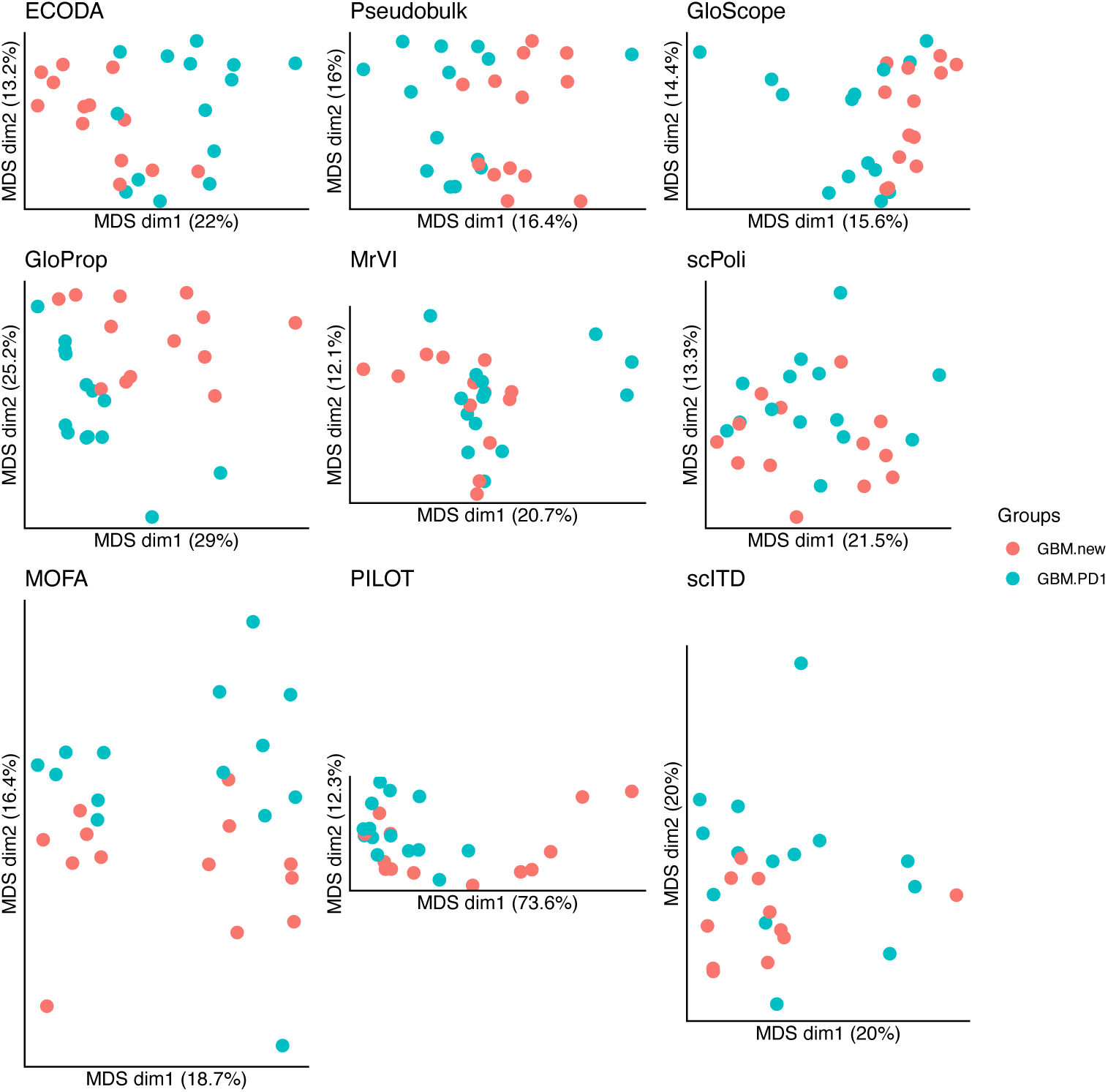
Classical Multidimensional Scaling (MDS) plots of inter-sample distance matrix of the Lee dataset. The groups in the Lee primary glioblastoma dataset are samples from newly diagnosed patients (GBM.new) or patients after anti-PD1 treatment (GBM.PD1).

**Supplementary Figure 8:**
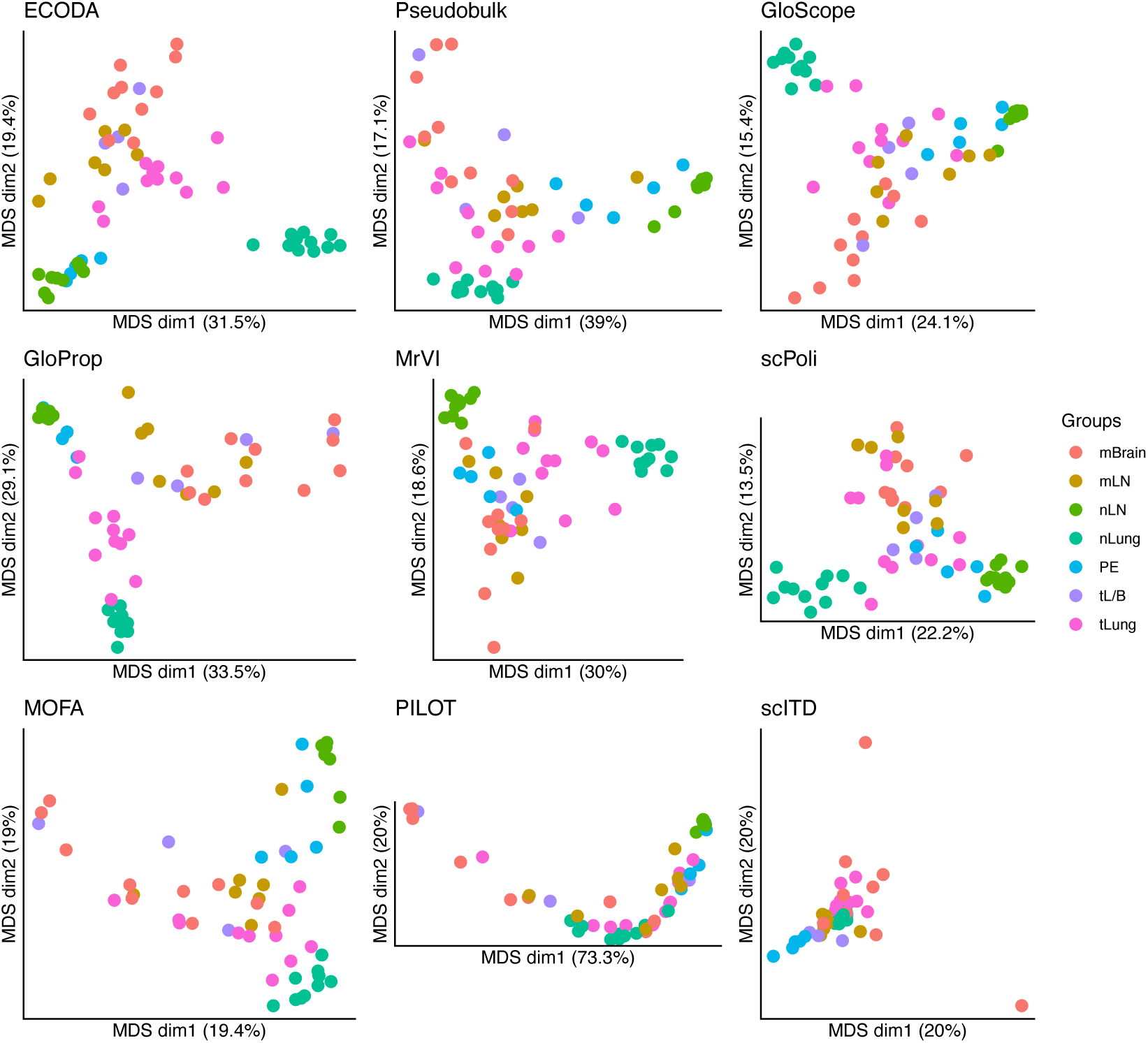
Classical Multidimensional Scaling (MDS) plots of inter-sample distance matrix of the Kim dataset. The groups in the Kim lung adenocarcinoma (LUAD) dataset are samples from normal Lung (nLung) or lymph node (nLN), pleural fluids (PE), tumor from lung at early stage (tLung) or late-stage biopsy (tL/B) and metastasis in brain (mBrain) and lymph node (mLN).

**Supplementary Figure 9:**
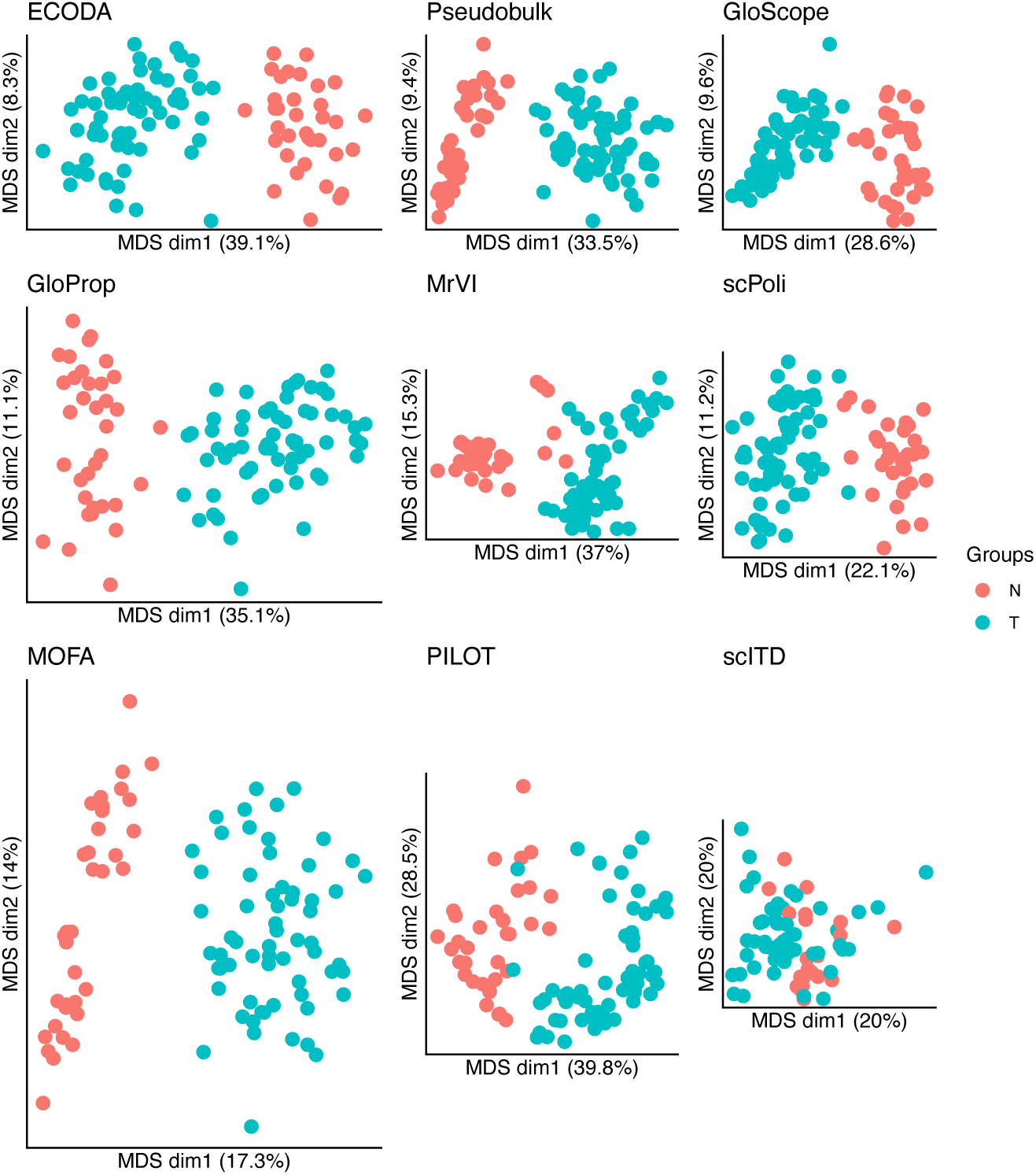
Classical Multidimensional Scaling (MDS) plots of inter-sample distance matrix of the Pelka dataset. The groups in the Pelka colon cancer dataset are samples from normal (N) or tumor (T) tissue.

**Supplementary Figure 10:**
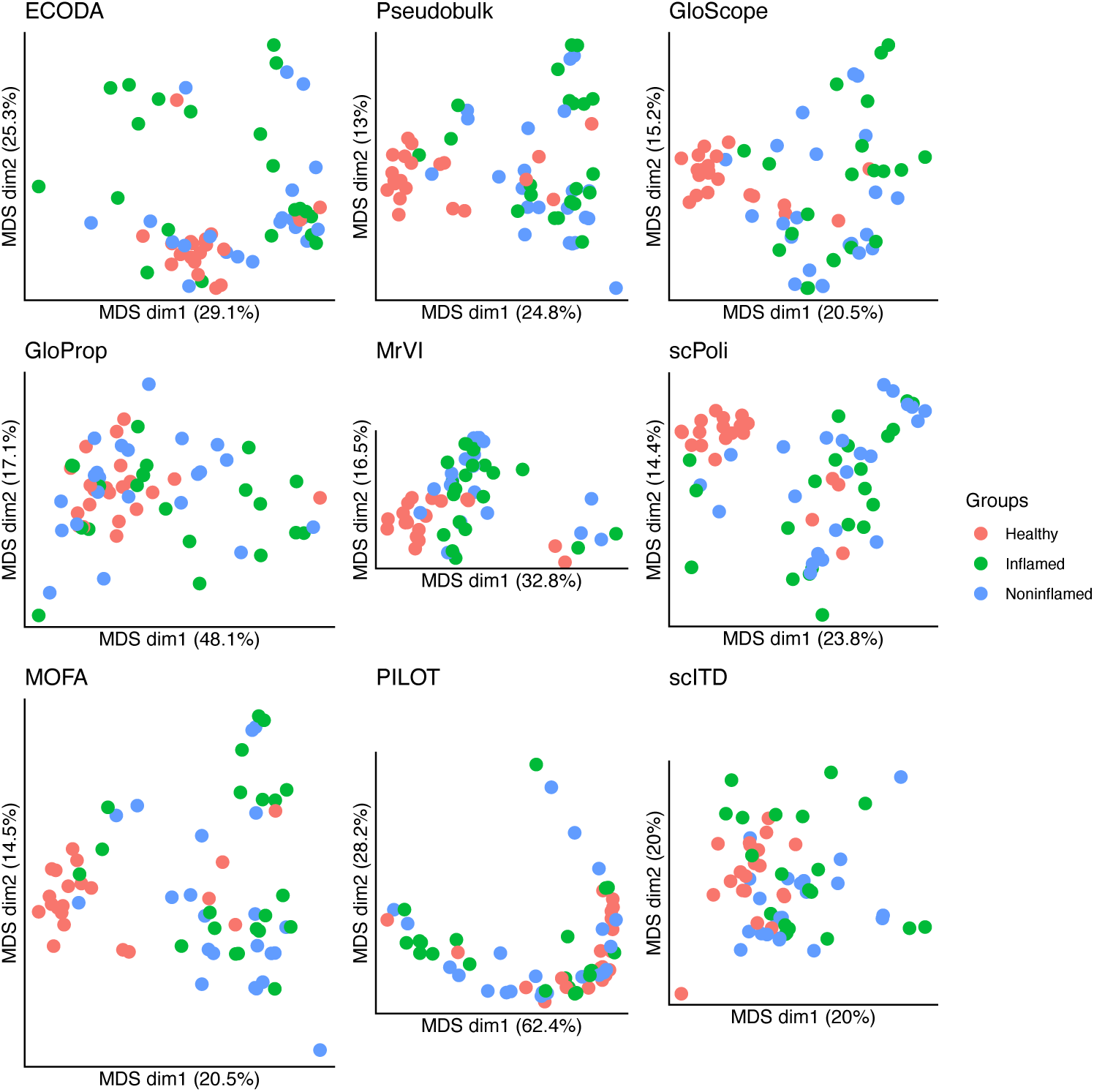
Classical Multidimensional Scaling (MDS) plots of inter-sample distance matrix of the Smillie dataset. The groups in the Smillie ulcerative colitis dataset are immune cell samples from lamina propria of either healthy controls, or patient samples from inflamed or adjacent non-inflamed colon.

**Supplementary Figure 11:**
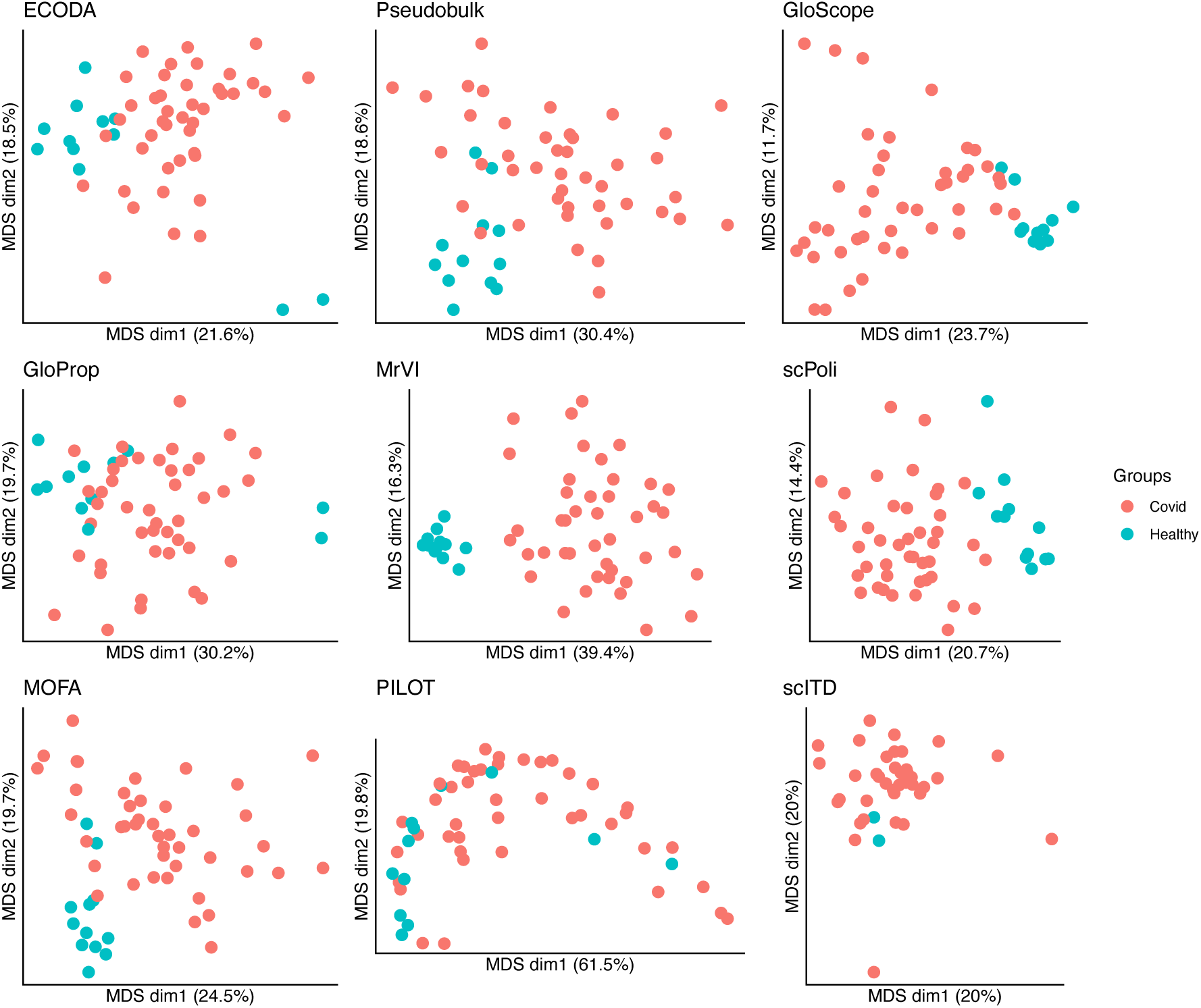
Classical Multidimensional Scaling (MDS) plots of inter-sample distance matrix of the Stephenson dataset. The groups in the Stephenson COVID-19 dataset are from healthy controls or acute COVID-19-infected people.

**Supplementary Figure 12:**
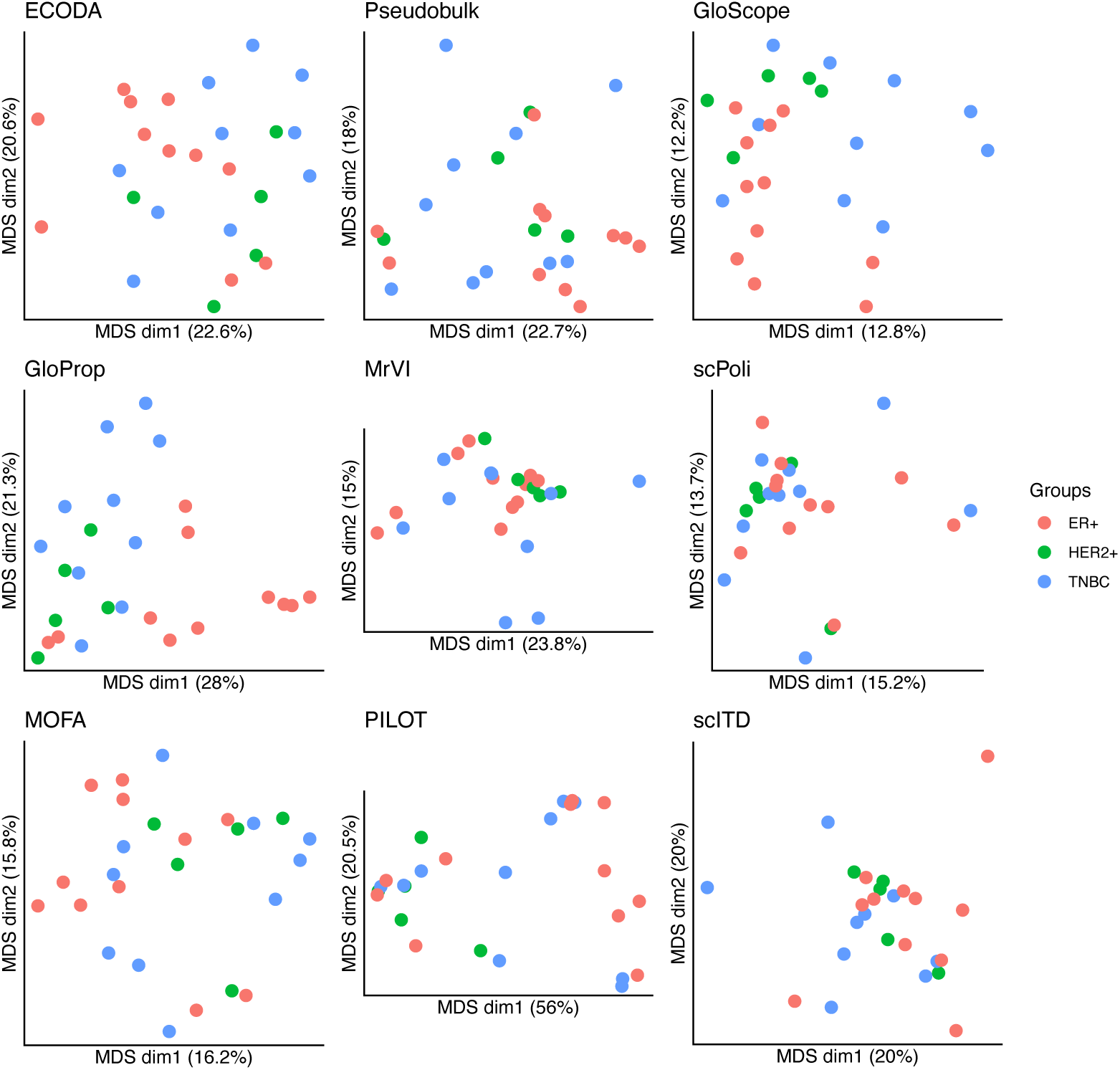
Classical Multidimensional Scaling (MDS) plots of inter-sample distance matrix of the Wu dataset. The groups in the Wu breast cancer dataset are samples from patients with different cancer subtypes (ER+, HER2+ or triple-negative breast cancer (TNBC)).

**Supplementary Figure 13:**
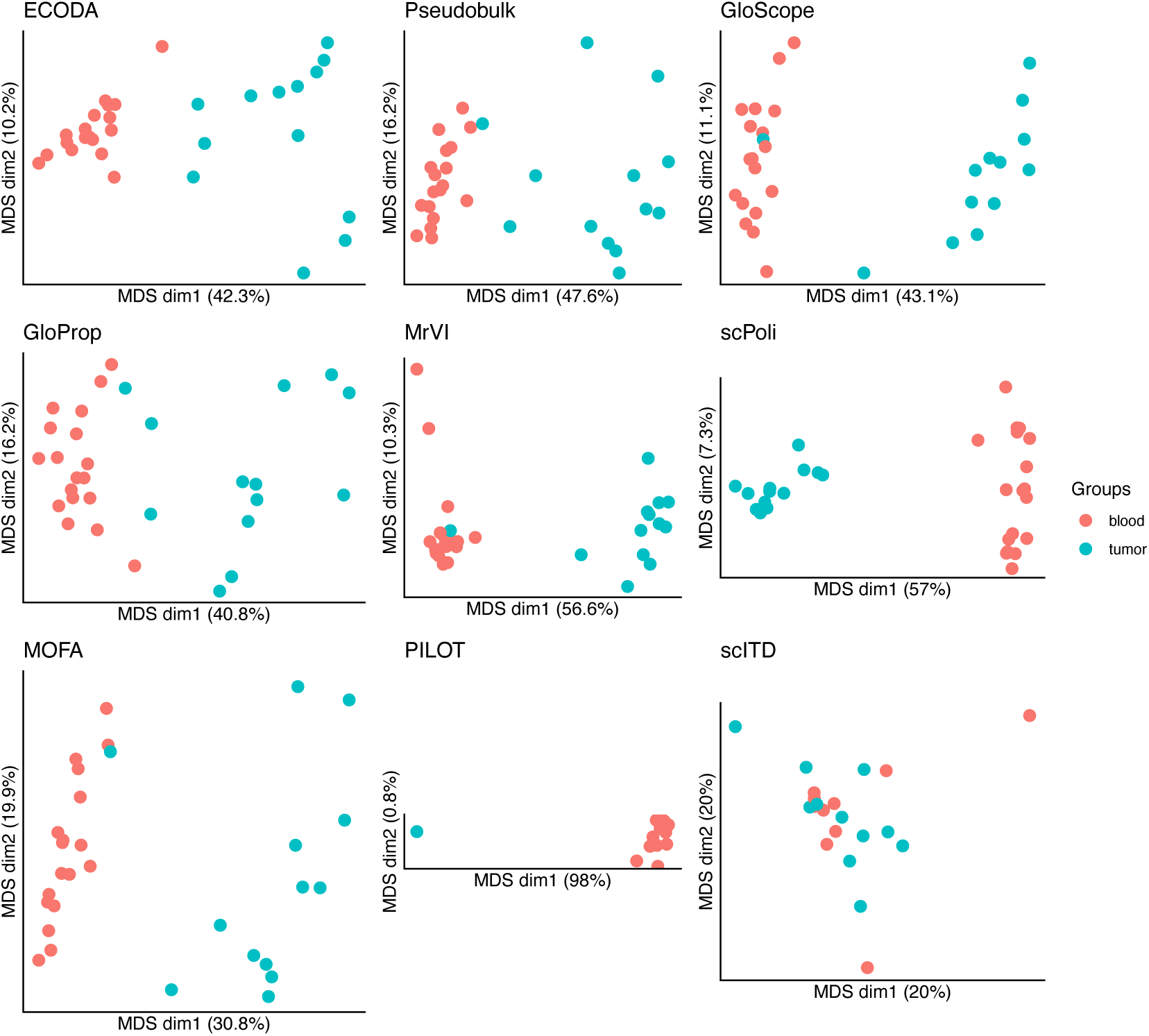
Classical Multidimensional Scaling (MDS) plots of inter-sample distance matrix of the Zhang dataset. The groups in the Zhang breast cancer dataset are blood or tumor samples.

**Supplementary Figure 14:**
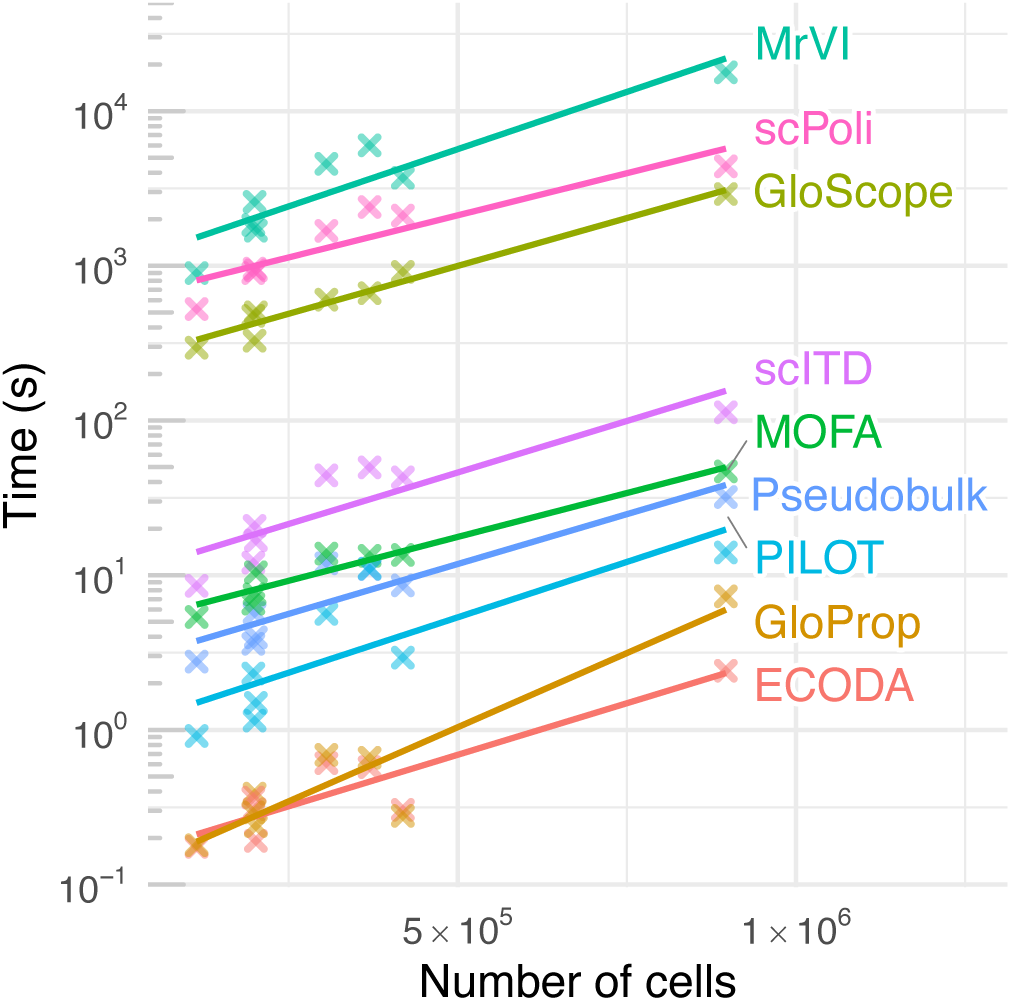
Method execution times. Each cross represents one dataset and how long the respective method took to process it. Lines show linear regression fit per method. Note: runtimes on the y-axis are shown on a log scale.

**Supplementary Figure 15:**
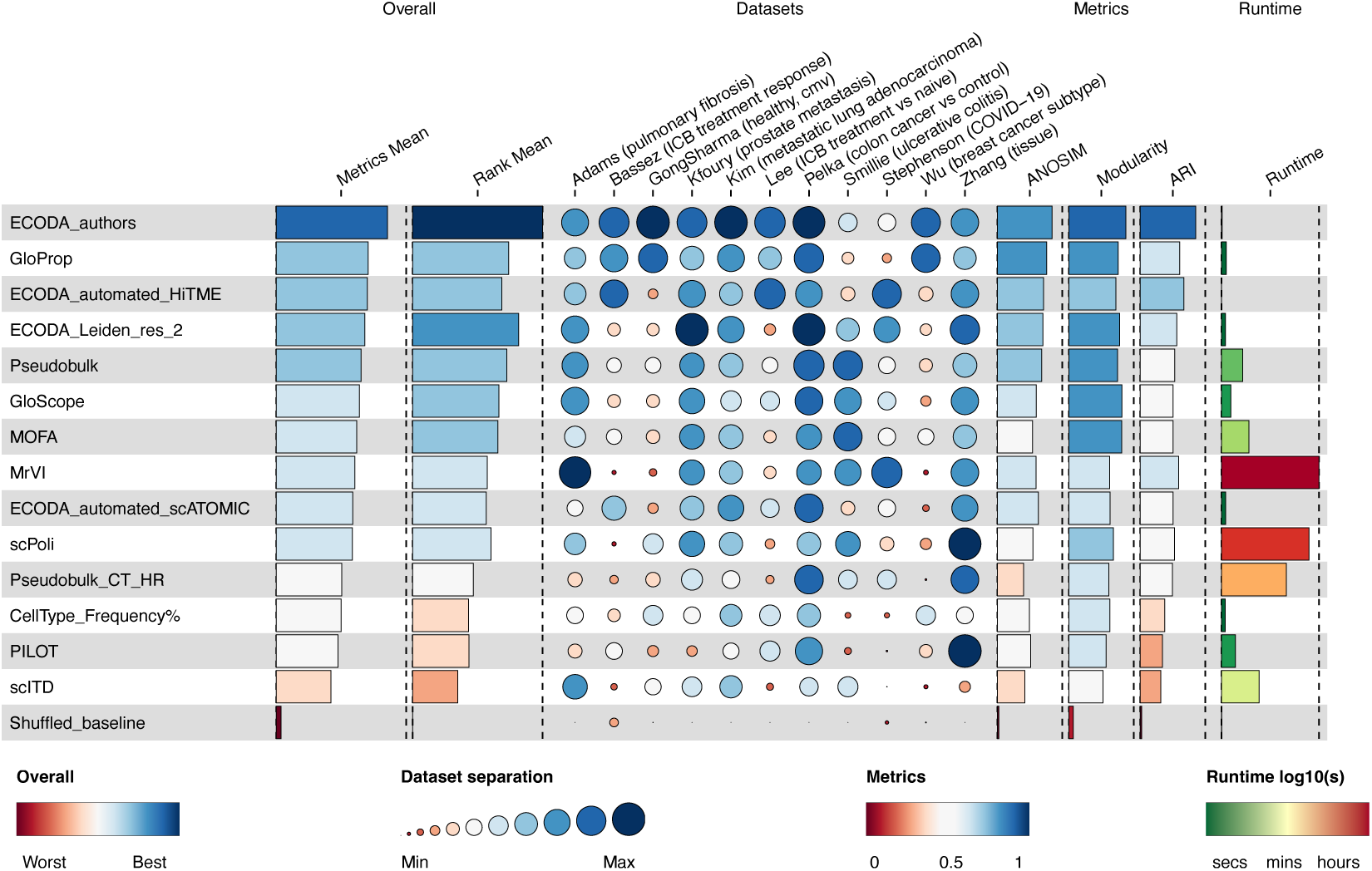
Extended benchmark of scRNA-seq sample representation methods for patient stratification (related to Figure 2). Representation methods are sorted by the average of the three separation metrics: Analysis of Similarity (ANOSIM), graph Modularity and Adjusted Rand Index (ARI) (“Metrics Mean”). Additionally, methods were ranked per dataset per metric and their rank geometric mean calculated, shown in the “Rank Mean” column. The dots display average metric scores per dataset. Separation metrics were min-max-scaled across the tested methods per dataset to range from [0, 1], to give equal weights to each dataset. ECODA is based on cell type labels annotated by authors (ECODA_authors), automated software tools (ECODA_automated_HiTME and ECODA_automated_scATOMIC) or unsupervised clustering of cells (e.g. using Leiden clustering with a resolution parameter of 2, “ECODA_Leiden_res_2”), see Methods. CellType_Frequency% represents the non-log-ratio-transformed cell-type proportions in percent. Pseudobulk_CT_HR refers to the average distance matrix across all high-resolution (HR) cell type pseudobulks. “Shuffled_baseline” is a negative control ECODA sample representation, where sample labels have been shuffled.

**Supplementary Figure 16:**
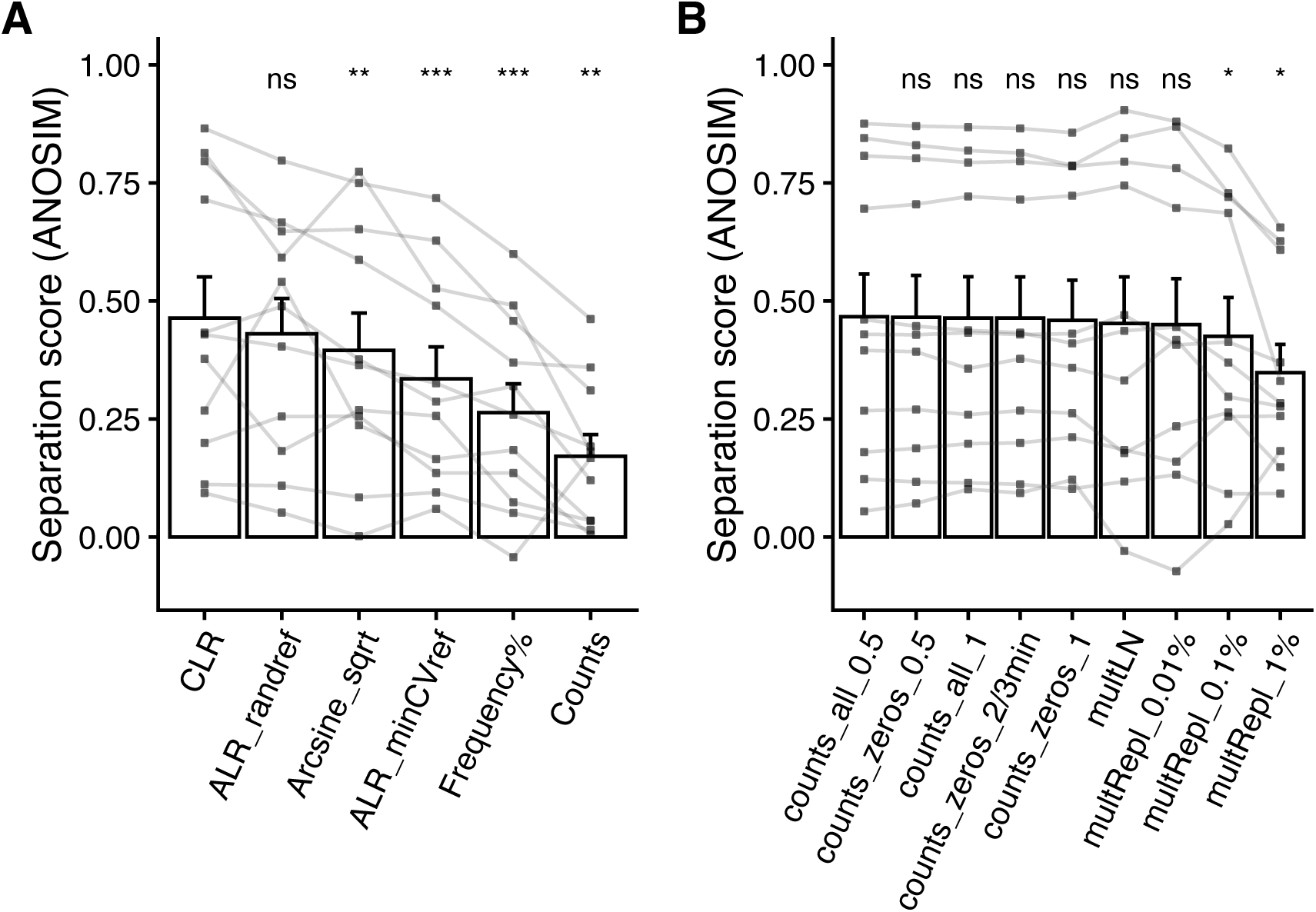
Effect of compositional data transformations for sample separation. A: Impact of normalization methods for cell-type compositional data on biological group separation. Centered Log-Ratio (CLR) transformation achieves significantly higher ANOSIM scores compared to relative frequencies (percent), arcsine square-root, and raw counts. B: Effect of zero-handling strategies on CLR-based separation. Bars represent the mean; whiskers indicate the standard error of the mean. Individual points represent specific datasets connected across methods. Asterisks indicate statistical significance from a paired Wilcoxon test against the CLR or counts_all_0.5 reference (ns: not significant, *: *p* < 0.05, **: *p* < 0.01, ***: *p* < 0.001), respectively.

**Supplementary Figure 17:**
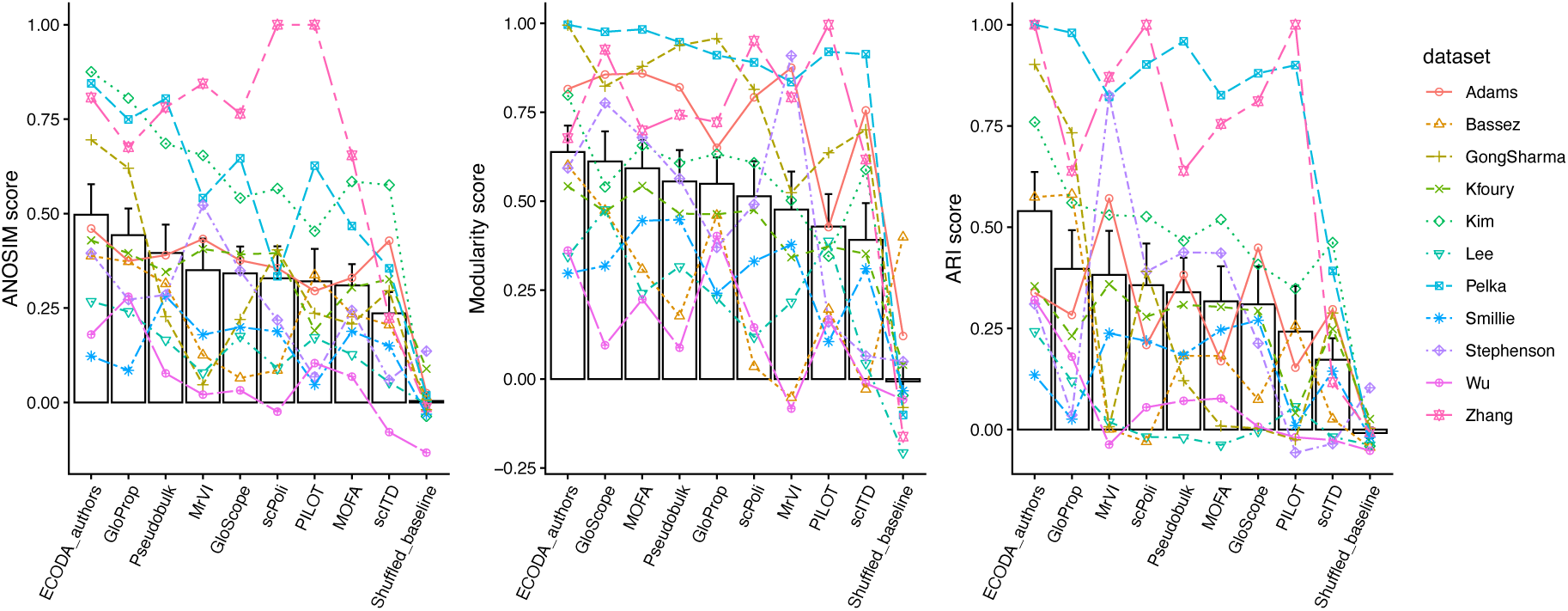
Unscaled performance metrics by dataset. Bars represent the mean, whiskers the standard error of the mean.

**Supplementary Figure 18:**
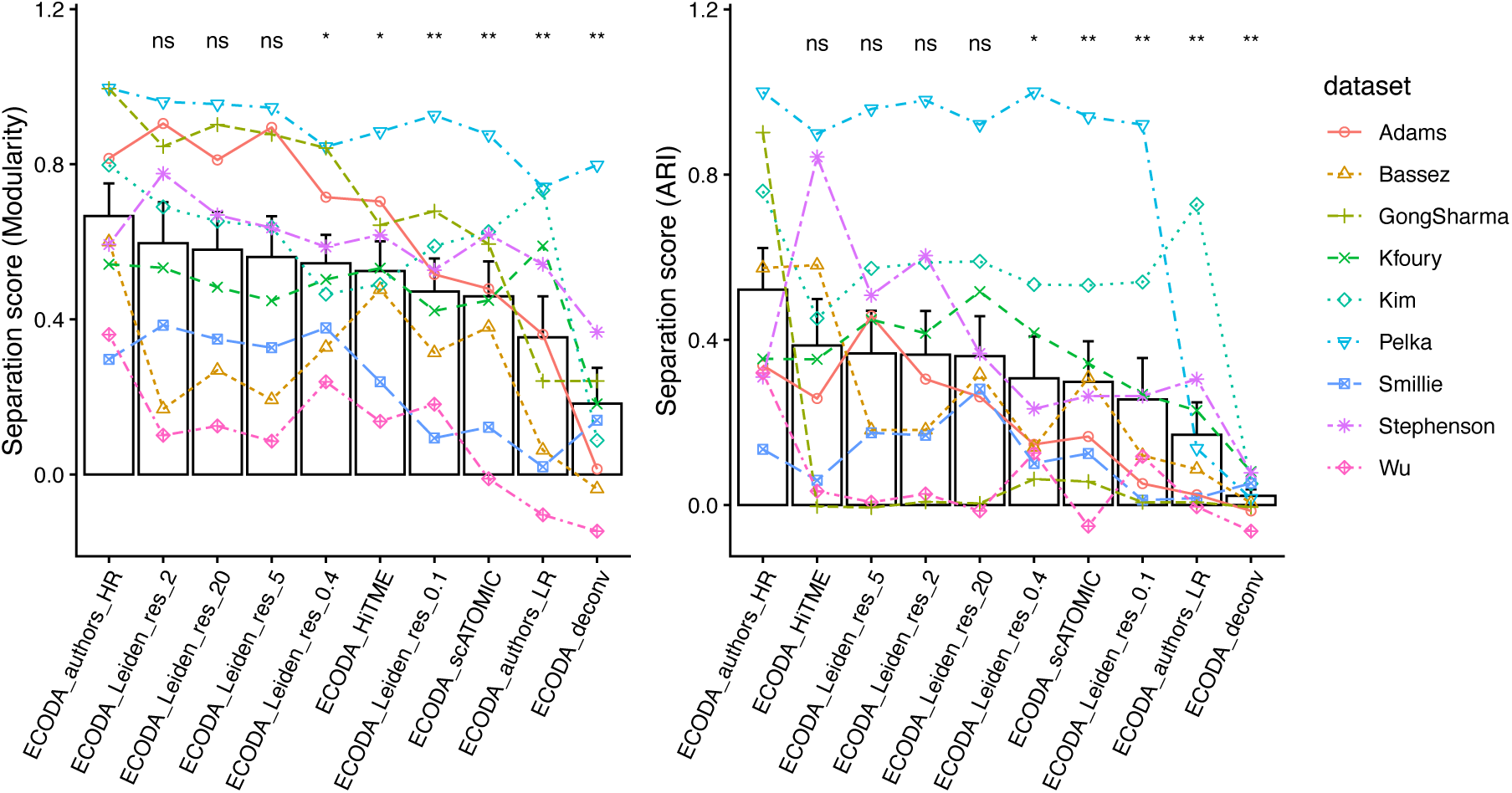
Robustness of ECODA performance across cell type annotation strategies evaluated by Modularity and Adjusted Rand Index. Comparison of Modularity (left) and Adjusted Rand Index (right) separation scores for various ECODA annotation strategies across ten biological conditions in eight datasets. Strategies evaluated include high-resolution (HR) author annotation, unsupervised Leiden clustering at multiple resolutions (0.1, 0.4, 2, 5, and 20), automated pipelines (HiTME, scATOMIC), low-resolution (LR) author annotation, and pseudobulk deconvolution. Asterisks indicate statistical significance from a paired Wilcoxon test against ECODA_authors_HR (ns: not significant, *: *p* < 0.05, **: *p* < 0.01).

**Supplementary Figure 19:**
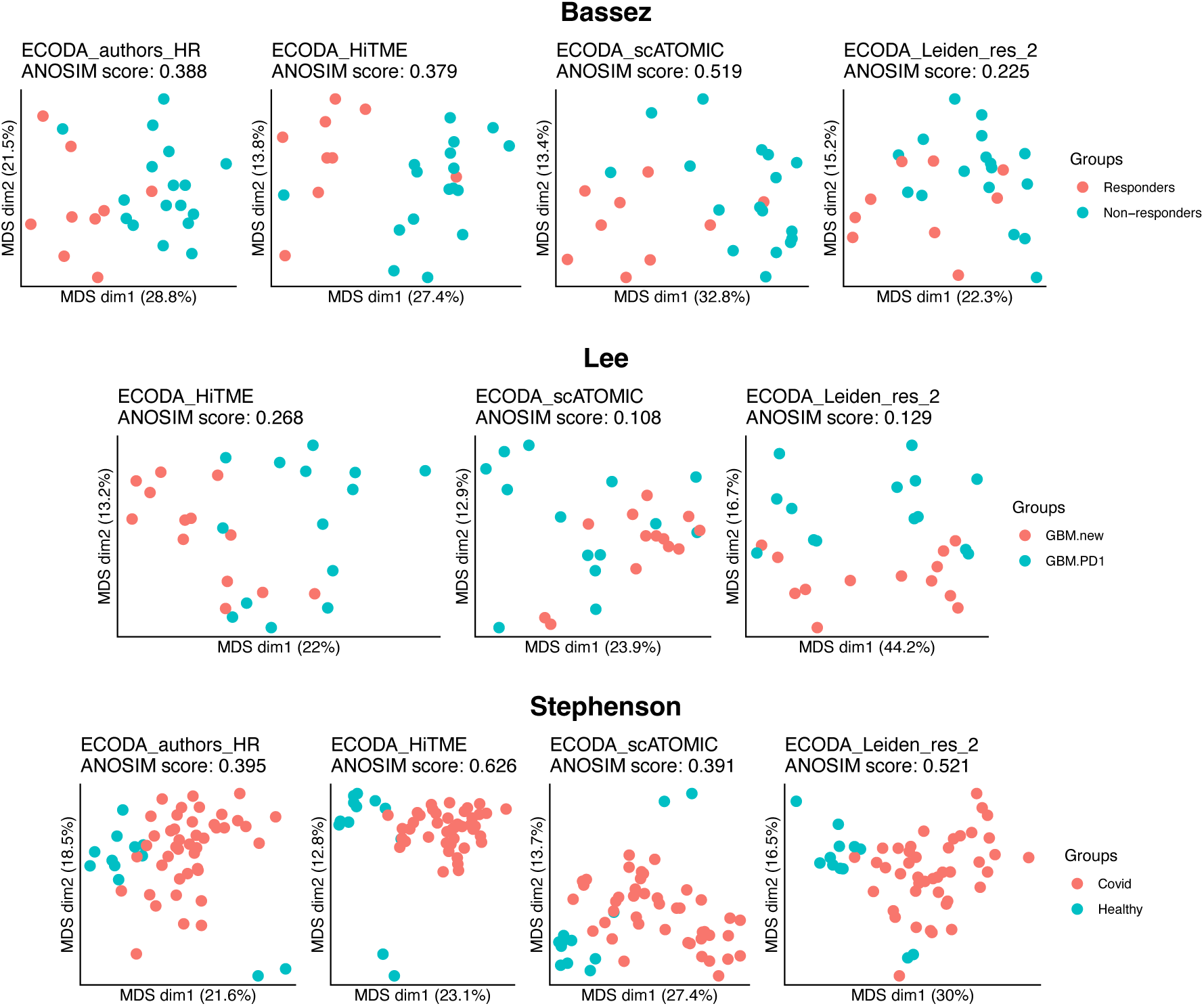
Examples where automated annotations outperform unsupervised clustering annotation. Shown are Classical Multidimensional Scaling (MDS) plots for the Bassez (top), Lee (middle) and Stephenson (bottom) datasets. Note: for the Lee dataset, no high-granularity author labels were available.

**Supplementary Figure 20:**
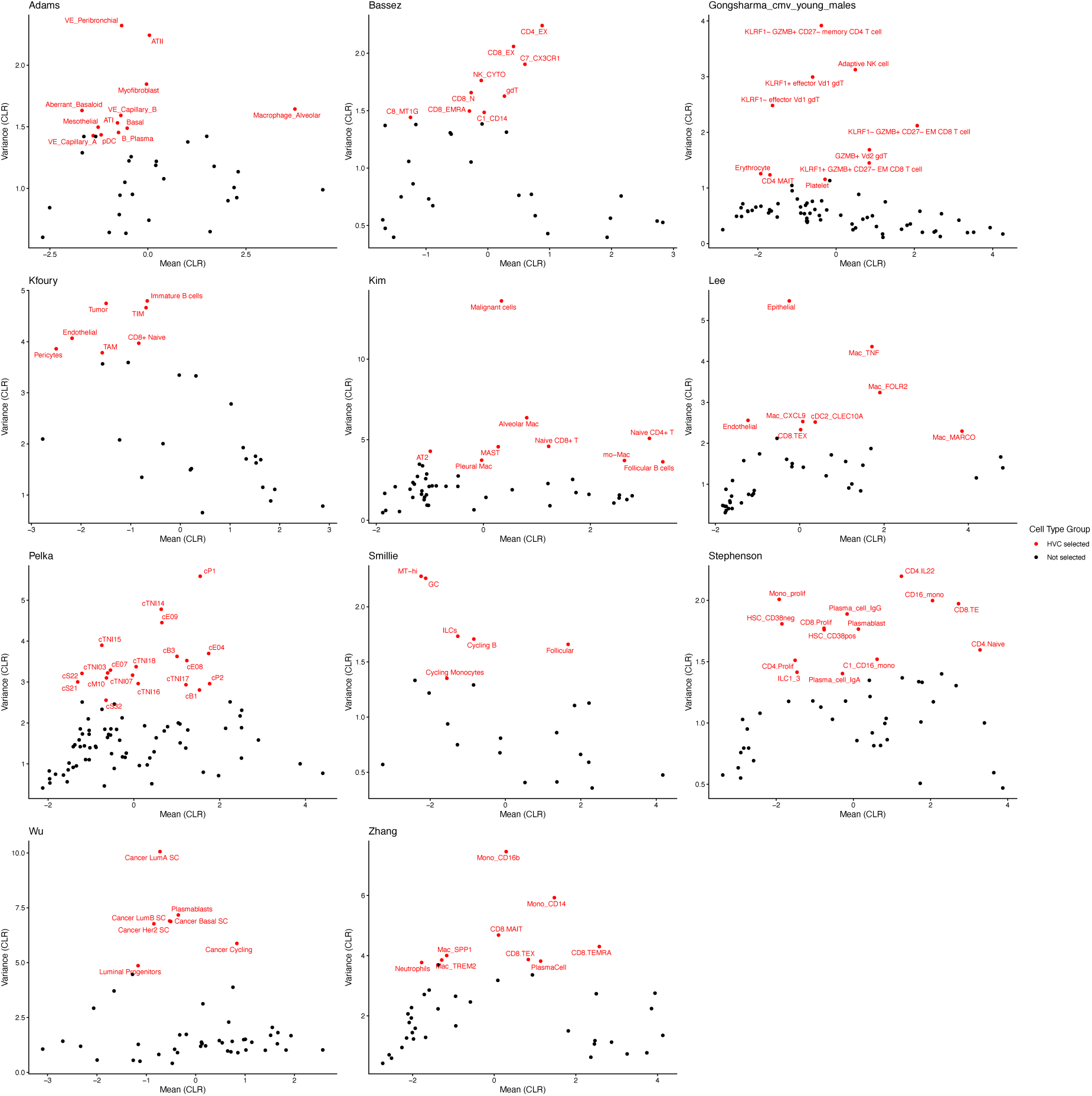
Mean-variance plots showing the most highly variable cell types per dataset. The mean-variance plots show the cell type abundances after centered log-ratio (CLR) transformation, with the mean (CLR) abundance on the x-axis and the variance (CLR) on the y-axis. The most highly variable cell types explaining 40% of the variance per dataset are highlighted in red (“HVC selected”).

**Supplementary Figure 21:**
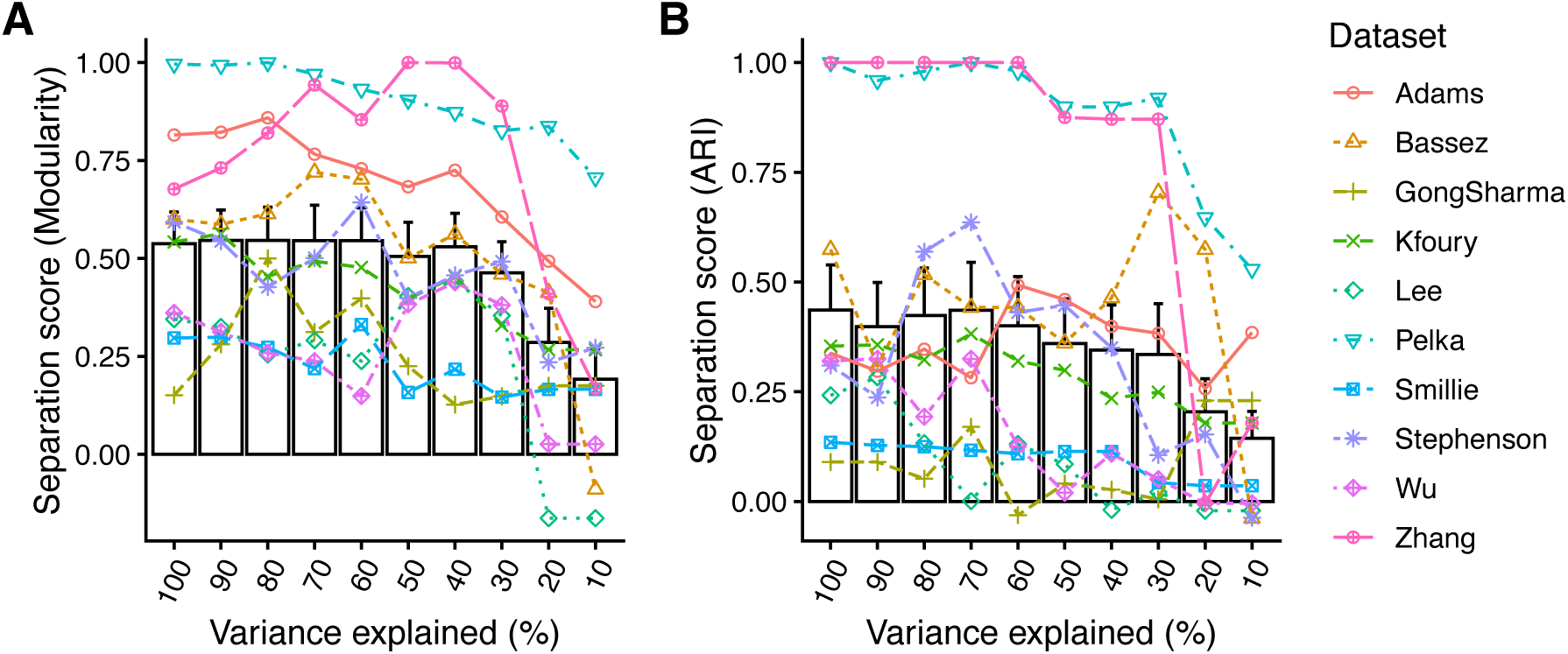
Variance explained by Highly Variable Cell Types (HVC). Modularity (A) and ARI (B) scores, when using only the HVCs explaining the variance indicated on the x-axis.

**Supplementary Figure 22:**
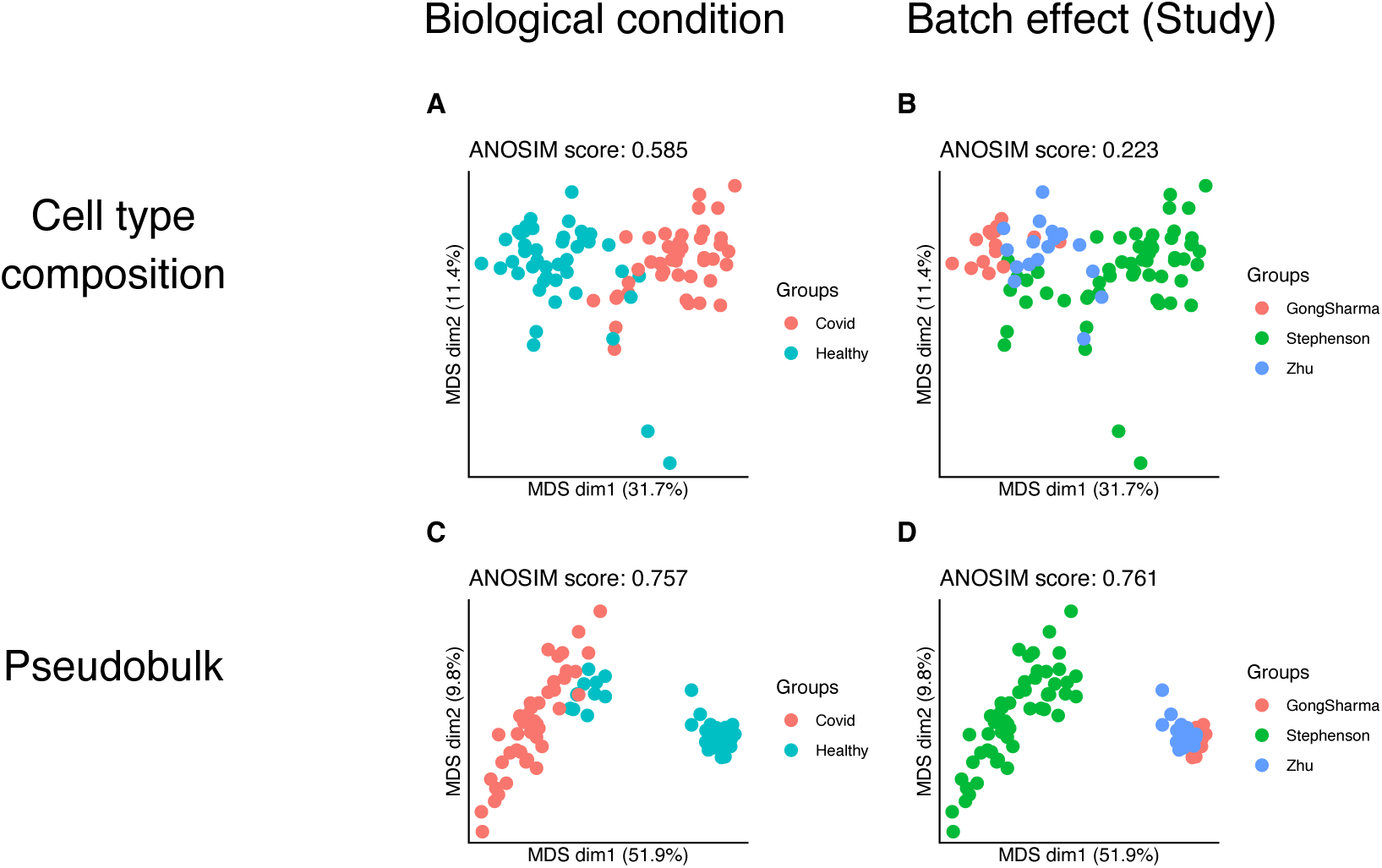
Comparison of biological signal and sequencing-related batch effects in a dataset comprised of data from multiple studies. Classical multidimensional scaling (MDS) plots illustrating the separation of biological groups and technical batches across two data representations. (A, B) Cell-type compositional embeddings (ECODA). (C, D) Pseudobulk gene expression embeddings. Left columns (A, C) are colored by biological condition (acute COVID infected, healthy control). Right columns (B, D) are colored by study of origin. Separation scores represent the mean of ANOSIM, modularity, and ARI metrics. Note that while pseudobulk displays higher technical clustering based on study of origin, cell type composition remains more robust to batch effect, preserving a higher relative biological signal (score: 0.585) compared to batch effect separation (score 0.223), while pseudobulk shows much stronger batch effect (score: 0.761), slightly higher than biological separation (score: 0.757). The Gong & Sharma cohort (180 samples) was subset to 15 randomly selected samples to balance the different datasets.

